# An immunocompetent murine model of virus-elicited liver fibrosis and hepatocellular carcinoma

**DOI:** 10.1101/2025.09.19.677076

**Authors:** Mariana Nogueira Batista, Juliano Bordignon, Ana Luiza Pamplona Mosimann, Tesia Bobrowski, Hsuan-An Chen, Gabriel Tobin-Xet, Erika A. Barrall, Nataliya Prokhnevska, Abishek Balachandra Vaidya, Tyler Lewy, Kenneth H. Dinnon, Leon Louis Seifert, Briana Zeck, Corrine Quirk, Yu-Jui Ho, Aveline Filiol, Raphael Wolfisberg, Caroline Jiang, Bruno Cogliati, Luis Chiriboga, Neil Theise, Margaret R. MacDonald, Alice Kamphorst, Troels K. H. Scheel, Timothy P. Sheahan, Eva Billerbeck, Scott Lowe, Brad R. Rosenberg, Charles M. Rice

## Abstract

Hepatocellular carcinoma (HCC) is the third deadliest cancer worldwide. Over 75% of HCC cases are associated with chronic viral infections. Mechanistic studies and preclinical therapeutic development for virus-associated HCC have been limited by a paucity of small animal models of chronic hepatotropic virus infection that faithfully recapitulate human disease. Here we demonstrate the induction of chronic hepatitis, progressive liver fibrosis, and HCC in immunocompetent laboratory mice upon chronic viral infection with Norway rat hepacivirus (NrHV) - a virus closely related to hepatitis C virus (HCV). NrHV-elicited tumors resemble HCV-associated tumors and liver transcriptome analyses reveal numerous similarities between chronic NrHV and HCV. These findings establish an experimentally tractable, physiologically relevant, and immunocompetent mouse model of virus-elicited progressive liver fibrosis and oncogenesis.

---

Hepatocellular carcinoma (HCC) is the most common form of liver cancer, and its high lethality makes HCC the third leading cause of cancer-associated mortality worldwide (*1*). Although new therapeutic modalities are constantly being explored, HCC incidence and mortality have increased over the past decade, and gaps in our understanding of pathogenesis and tumor vulnerabilities persist (*1, 2*). The oncogenic process of HCC unfolds over decades in humans, presenting challenges for longitudinal analyses. Furthermore, the remarkable heterogeneity of tumors within and among different etiologies hinders proof-of-concept studies (*2*). Chronic infections with hepatitis B virus (HBV) and hepatitis C virus (HCV) are among the leading causes of HCC worldwide. Despite the availability of highly effective direct-acting antivirals (DAAs), HCV remains the dominant cause of HCC in the United States, Western Europe, North Africa, the Middle East, and Japan (*2*). The extent of liver damage tolerated by the host, potential of damage reversion, and its contribution to HCC development remain critical, unresolved research questions (*3, 4*) but viral hepatocarcinogenesis remains difficult to model.

HCV is a hepacivirus that infects humans. Historically, the only experimental animal models capable of supporting HCV infection are chimpanzees and treeshrews (*5*). However, chimpanzee studies are limited by ethical concerns, high genetic variability, low HCC incidence following infection, and high costs. Treeshrews present challenges such as genetic variability and a lack of species-compatible research tools (*5*). Other non-human primates, including rhesus and cynomolgus macaques, do not support productive HCV infection (*5*). Due in part to the broad availability of research tools, established preclinical workflows, and relatively low cost, mice are a preferred animal model for pathogenesis and immunological studies. However, wild-type murine hepatocytes are not susceptible to HCV infection. Strategies developed by our group and others to study HCV-associated pathogenesis in murine models include virus adaptation, “humanization” of viral entry receptors, and the engineering of immunodeficient mice harboring xenotransplanted human liver tissue (*5*). To date, however, none of these models recapitulate advanced hepacivirus-associated disease such as HCC (reviewed in (*5*)). In an innovative alternative approach, hepaciviruses identified in other animal species can be used as surrogate models for HCV (*5–7*), with the possibility that related viruses within the genus cause similar disease in their respective (and related) hosts.

We have developed and characterized the first immunocompetent laboratory mouse model supporting chronic hepacivirus infection using Norway rat hepacivirus (NrHV), a natural pathogen of New York City rats (*8, 9*). NrHV shares numerous similarities with HCV, including strict hepatotropism, genome organization, replication mechanisms, and host entry factor engagement (*8, 10–16*). Initial studies demonstrated that acute NrHV infection in laboratory mice and rats elicited immune responses similar to those observed in acute HCV infection of chimpanzees and humans (*8, 10, 17*). In addition, NrHV clearance is mediated by CD8 T cells and is dependent on CD4 T cells, similar to what is observed in humans(*8, 18*). Transient depletion of CD4 T cell enables persistent hepacivirus infection in both chimpanzees (HCV) and mice (NrHV); importantly, persistent infection is sustained upon CD4 T cell recovery (7, 13–16). Furthermore, as in chronic HCV infections in humans, chronic NrHV infection in mice is marked by an increased frequency of regulatory T cells (Tregs) and CD8 T cell exhaustion (*8*). Given that the NrHV mouse model has been established as a surrogate for acute HCV infection, we hypothesized that extended chronic NrHV infection in wild-type mice would lead to progressive liver disease modeling chronic HCV-associated pathology. Herein, we characterize the progression of chronic NrHV infection in immunocompetent C57BL/6J (B6) mice over an 18-month period and demonstrate that this model recapitulates key features of chronic hepacivirus pathogenesis in humans, including hepatocarcinogenesis.

## Mouse-adapted NrHV establishes chronic infection in wildtype C57BL/6J mice and is controlled earlier and more efficiently in females versus males

We have previously shown that transient depletion of CD4 T cells prior to NrHV infection of wild-type (WT) B6 mice results in chronic infection, with viral persistence lasting at least 7 months despite full recovery of CD4 T cells by 1-month post-infection (m.p.i) (*8*). To evaluate the long-term consequences of chronic NrHV infection, we infected 10–12-week-old B6 mice with NrHV-B_SLIS_ (mouse-adapted NrHV) (*11*) 4 days after CD4 T cell depletion, and monitored viremia and associated clinical signs for 18 months. A transiently CD4-depleted mock-infected group was followed in parallel as control (*Cohort1*; n=15 mock vs n=60 NrHV) (**Fig. 1A**). NrHV infection persisted for up to 18 months in approximately 70% of infected mice (**Fig. 1B, fig. S1**). NrHV viremia peaked at one m.p.i, reaching 10^8^ genome equivalents (GE)/mL prior to a gradual 100-fold decrease over the next 5 months that eventually stabilized. Female patients are more likely to clear HCV infection than males (*19*), and similarly, we found that female mice were more likely than males to eventually resolve chronic NrHV infection than males. Viremia was significantly lower in females in the first month post infection (p<0.0001), but there was no statistically significant difference in viral genome RNA levels between infected males and females at later time points (**fig. S1**).

**Fig. 1.**
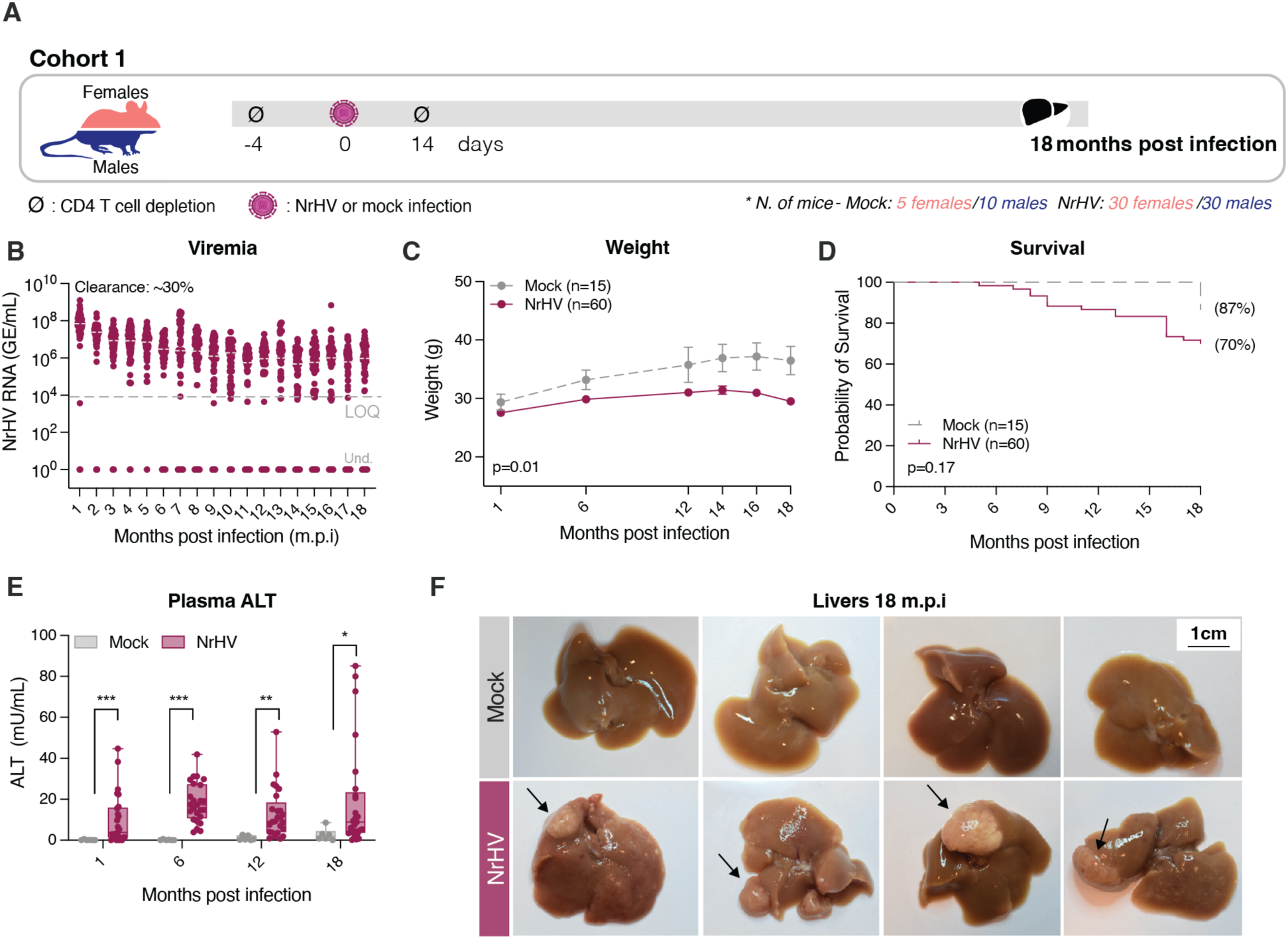
Chronic NrHV infection leads to liver damage and tumors in wild-type mice. **(A)** Experiment schematic: wild-type C57BL/6J mice (Cohort 1 n=75) were depleted for CD4 T cells 4 days prior and 14 days post infection. At day zero mice were either infected with 10^4^ genome equivalents (GE) of NrHV (n=60) or mock-infected (n=15); number of males are indicated in blue and female in red. Blood was collected monthly and organs harvested at 18 months post infection (m.p.i). **(B)** Viremia in NrHV-infected mice during the 18 months of experimentation. Und. = undetectable viremia. **(C)** Mice body weight in NrHV-infected (maroon) and mock-infected (gray) over time. Statistics: Mixed-effects model for repeated measures, n=75, ***p=0.0001 NrHV vs mock over time. **(D)** Kaplan-Meier survival analysis. Maroon: NrHV-infected, Gray: mock controls. Percentage indicates probability of survival in each group. Statistics: Log-rank (Mantel-Cox) test, n=75, p=0.17. **(E)** Alanine aminotransferase (ALT) activity measured in plasma of mice over time. Statistics: Mann-Whitney test Time 1: p=0.0032, Time 7: p=0.0002, Time 12: p=0.0009, Time 18: p=0.0153 NrHV vs Mock, alpha 0.05, Bonferroni adjusted alpha 0.0125 (n=31). **(F)** Liver gross view in chronic NrHV-infected and mock-infected mice. Tumors indicated by black arrows.

## Chronic NrHV infection results in elevated plasma ALT levels and liver tumors

Chronic NrHV infection impacted body weight over the course of infection, resulting in trends of delayed increases in body weight in females and statistically significant delayed increases in males compared to mock controls (**Fig. 1C, fig.S2A**). Although most mice survived the entire 18-month study, we observed a trend towards higher mortality rate in NrHV-infected mice compared to mock-infected controls, that was significant in male mice when analyzed separately from female mice (**Fig. 1D, fig. S2B**). HCV infection often causes subclinical liver injury, typically assessed by serum or plasma alanine transaminase (ALT) levels. NrHV infected animals exhibited significantly elevated ALT levels compared to mock controls (1 m.p.i: p=0.0032; 6m.p.i: p=0.0002, 12 m.p.i: p=0.0009, 18 m.p.i: p=0.0153) (**Fig. 1E, fig. S2C**). Of note, relative to mock-infected controls, males exhibited significantly higher ALT levels than females at 1 and 6 m.p.i., but not at 12 and 18 m.p.i. (1 m.p.i: p=0.02; 6 m.p.i: p<0.0001). This suggested that males experienced more pronounced liver damage at earlier stages of chronic infection. Liver/body weight ratio was increased in infected males but in females the analysis suggested no change 18 m.p.i (males: p=0.009; females: 0.06) and spleen/body ratio sizes tended to be increased in both sexes in the infected group but not significantly (**fig. S2D-E**). At 18 m.p.i, liver gross view evaluation revealed proliferative lesions in 91% of all NrHV-infected mice consistent with hepacivirus-associated HCC (**Fig. 1F, table S1 and table S2**).

## Chronic NrHV infection causes hepatitis and progressive liver fibrosis in mice

To confirm our findings on liver tumor development and study the progression of liver disease during chronic NrHV infection in mice, we repeated our initial 18-month experiment with additional intermediate time points (*Cohort 2*; n=36 mock vs n=60 NrHV mice, **Fig. 2A**). Hepatitis, characterized by immune infiltration and hepatocyte necrosis, is a histopathological hallmark of chronic HCV, and is accompanied by portal lymphoid aggregates (LA) in about 60% of patients (*20*). Livers of chronic NrHV-infected mice exhibited high levels of inflammation with lobular hepatitis and portal LA. Comparable inflammation was not found in mock-infected controls (**Fig. 2B-C**). LA were present in 90% of the NrHV-infected mice at 18 m.p.i, with no significant difference in LA frequency between males and females at any time point (**fig. S3**). In addition to portal LA and lobular hepatitis, many of the histopathological features observed in livers from mice chronically infected with NrHV are also commonly observed in livers of HCV-infected patients, including steatosis, acidophil bodies, and Mallory-Denk bodies, indicating endoplasmic reticulum stress or misfolded proteins (**Fig. 2D**).

**Fig. 2.**
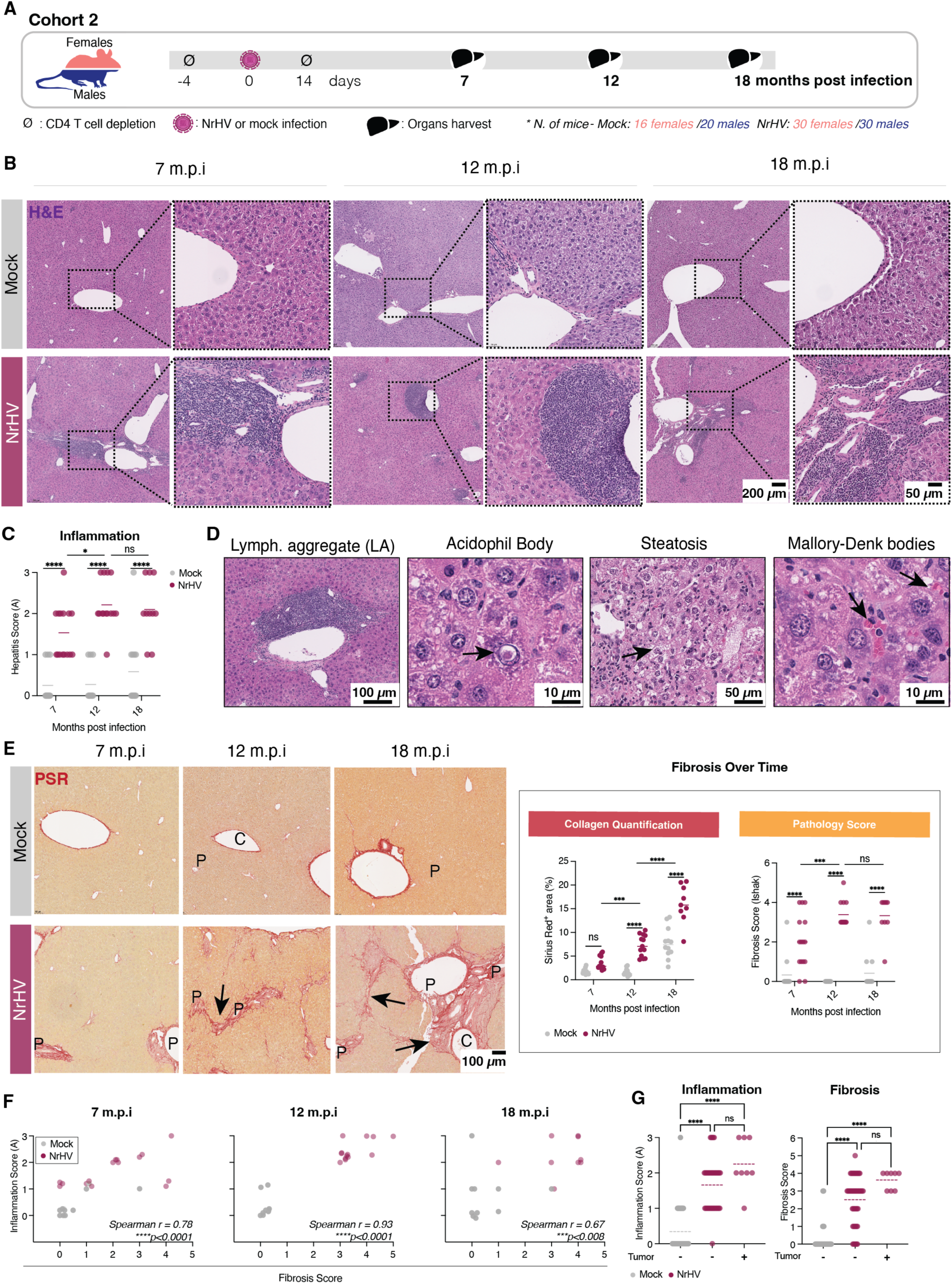
Chronic NrHV infection in mice recapitulates histopathological features of chronic HCV infection in humans. **(A)** Experimental scheme: Wild-type C57BL/6J mice (Cohort 2 n=96) were treated as in Cohort 1, blood was collected monthly and organs harvested at 7-, 12- and 18-months post infection (m.p.i); number of males are indicated in blue and females in salmon. **(B)** Hematoxylin/ Eosin (H&E) staining of representative livers 7- (n=21), 12- (n=24) and 18-months (n=21) post NrHV or mock infection. Statistics: Two-way ANOVA, time: p=0.04*, status: p<0.0001****, Sidák’s post-test p<0.0001 (n=72, alpha 0.05) for all comparisons. **(C)** Representative images of the main histopathological findings in chronic NrHV-infected mice that are comparable to those seen in chronic HCV-infected humans. **(D)** Picro Sirius Red (PSR) staining of representative livers 7-, 12- and 18-months post NrHV or mock infection followed by collagen quantification per area and pathology score. Collagen quantification - Statistics: Two-way ANOVA, time: p=0.02*, ns: non-significant; status: p<0.0001****, Sidák’s post-test p<0.0001 (n=72, alpha 0.05) for all comparisons. **(E)** Scatter plot for inflammation (A0=minimum activity/A3=maximum activity) and fibrosis (0=no fibrosis/6= cirrhosis). Each dot represents a single mouse liver. Statistics: Spearman correlation (n=72, alpha 0.05). P: portal tract; C: central vein. **(F)** Inflammation and Fibrosis Scores comparing livers from mice that did or did not develop tumors 18 months post infection. Statistics: Kruskal-Wallis test - Fibrosis score: p < 0.0001; Dunn’s post-hoc test adjusted for multiple comparisons Mock vs. NrHV HCC-: p < 0.0001****; Mock vs. NrHV HCC+: p < 0.0001****. Hepatitis score: p < 0.0001; Dunn’s post-hoc test adjusted for multiple comparisons: Mock vs. NrHV HCC-: p < 0.0001****, Mock vs. NrHV HCC+: p < 0.0001****, NrHV HCC- vs. NrHV HCC+: n.s. (not significant). For all panels gray: Mock-, maroon: NrHV-infected.

Liver fibrosis at various stages is observed in 85-98% of HCV patients. During the progression from periportal to bridging fibrosis, approximately 5-15% of chronic HCV cases eventually advance to cirrhosis (*21*). In livers from NrHV-infected mice, collagen accumulation increased over time, with evident portal tract expansion at 7 m.p.i., initial bridging fibrosis (portal-portal) or advanced bridging fibrosis (portal-central) at 12 m.p.i., and extensive collagen accumulation with early nodule formation at 18 m.p.i. (**Fig. 2E**). Blinded samples were scored by a board-certified pathologist using human histological scoring criteria for inflammation and fibrosis (METAVIR and Ishak, respectively) (**table S3**)(*22*). Inflammation and fibrosis scores were consistently higher in NrHV-infected mice in comparison to mock controls with no significant difference between infected males and females over time (**fig. S4**). Fibrosis and inflammation scores were variable from mouse to mouse but overall correlated, with the strongest correlation occurring 12 m.p.i. (**Fig. 2F**). Tumors were observed in association with various levels of liver inflammation but only in contexts of advanced fibrosis (**Fig. 2G**). Spontaneous viral clearance, independent of the time post-infection, was associated with reduced liver inflammation, suggesting that ongoing NrHV infection is responsible for liver inflammation and that this state is reversible (**fig. S5**). Fibrosis, but not inflammation scores were affected by time of clearance, (Inflammation: r=0.48, ns. Fibrosis: r=0.84, p<0.001) (**fig. S5B**), suggesting that reversion of fibrosis is time-sensitive. However, due to unpredictable viral clearance in a minority of infected animals, further studies are needed to clarify fibrosis dynamics following spontaneous or drug-mediated cure.

## Chronic hepacivirus infection results in similar gene expression changes in the livers of humans and mice

Given the histological similarities between chronic NrHV and HCV infection, we next characterized the gene expression changes elicited by chronic NrHV. Principal component analysis of RNA sequencing (RNA-Seq) on liver tissue comparing mock, NrHV and cleared NrHV showed that expression in liver samples from animals that spontaneously cleared virus more closely resembled gene expression from mock-rather than chronic NrHV-infected animals **Fig. 3A**) - reinforcing that NrHV-elicited changes in liver tissue are to some extent reversible upon viral clearance. Therefore, subsequent comparisons were conducted in chronic versus mock animals only. RNA-Seq from mice at 7, 12, and 18 m.p.i showed that expression patterns were generally consistent across the infection time course (1,856 genes increased, 362 genes decreased) (**Fig. 3B**). Gene set enrichment analysis (GSEA)(*23*) at each timepoint highlighted an increase of interferon, inflammatory and epithelial-mesenchymal transition (EMT) programs in NrHV-infected livers compared to mock-infected, along with a downregulation of prototypical liver functions such as lipid and bile acid metabolism (**Fig. 3C**).

**Fig. 3.**
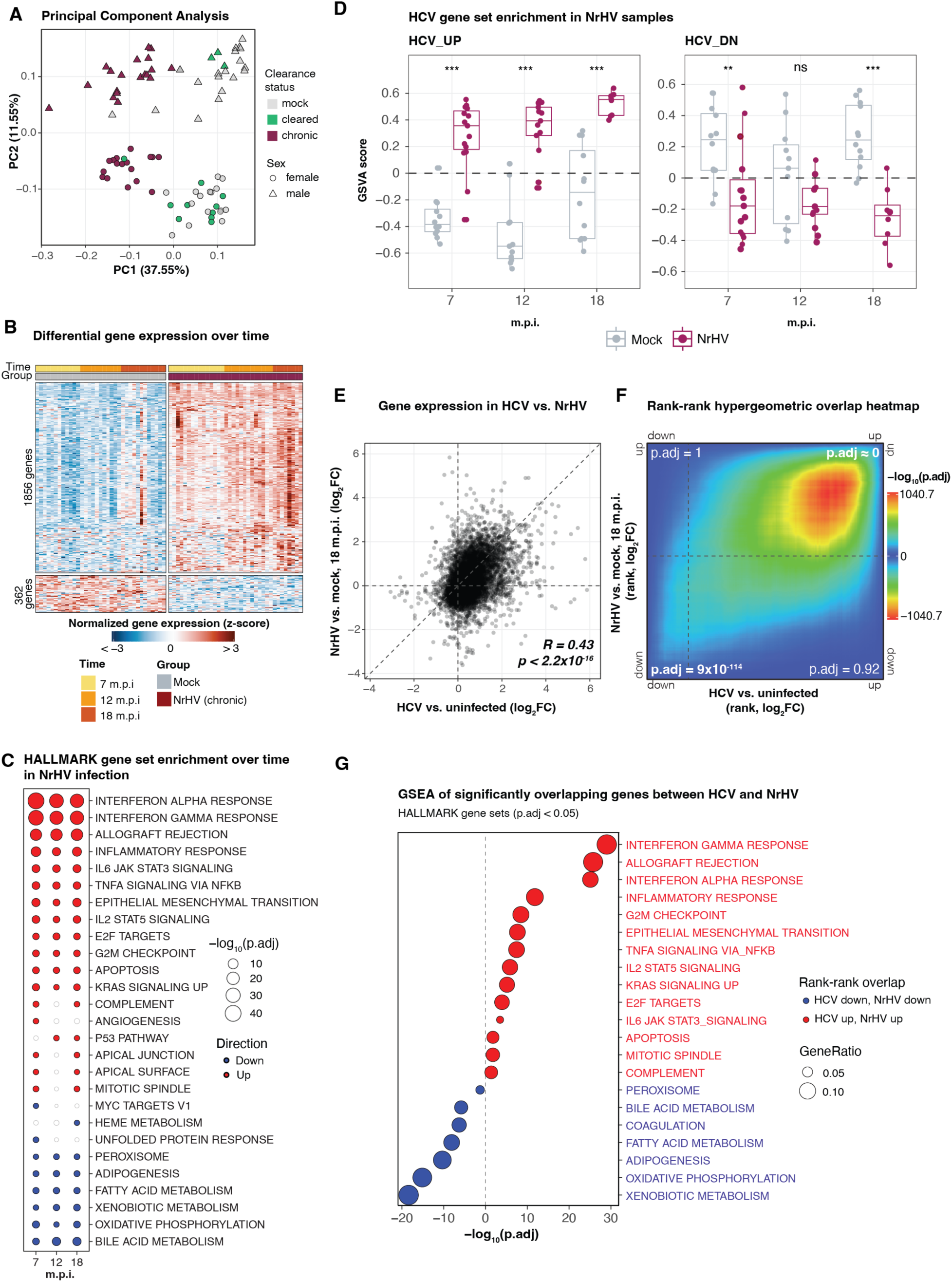
Chronic NrHV infection elicits gene expression changes associated with immune and liver metabolic functions highly similar to gene expression changes in chronic HCV. **(A)** Principal component analysis (3,000 most variable genes by RNA-Seq) of liver tissue from chronic NrHV- vs. mock-infected mice (cohort 2, all time points, including samples from mice that spontaneously cleared NrHV infection). **(B)** Heatmap (RNA-Seq) of differentially expressed genes (DEGs, p.adj <0.05 and log_2_FC >1 or <-1, any timepoint) in livers from NrHV- vs mock-infected mice 7 (n=30), 12 (n=27), and 18 (n=25) m.p.i. Across all time points there were 1,856 genes increased, 362 genes decreased. **(C)** Dot plot of HALLMARK gene sets significantly enriched (p.adj <0.05) at individual time points post infection in liver tissue from chronic NrHV- vs. mock-infected mice. Point size indicates significance; color indicates direction of enrichment (red = enriched in chronic infection; blue = enriched in mock infection). (**D**) GSVA analysis of HCV gene sets (HCV_UP and HCV_DOWN, *see Methods and Main* text) in chronic NrHV- and mock-infected liver tissue. Points represent samples from individual mice (mock N=35, NrHV N=37). Significance calculations performed with linear-model–based testing approach (with empirical-Bayes shrinkage): ***p < 0.001, ** p < 0.01, *p < 0.05, ns = not-significant (p >= 0.05). (**E**) Log2FC values for all one-to-one mouse-human orthologue genes in two independent RNA-Seq analyses: liver from chronic HCV infection versus uninfected (*data from Boldanova et al 2017 and Ramnath et al., 2018* (*24, 25*)) and liver from 18 m.p.i. chronic NrHV infection versus mock infection (*this study*). *R*, Pearson correlation coefficient. *Nnt* gene excluded as out of range (NrHV comparison, log_2_FC < −9). (**F**) Overlap map for rank-rank hypergeometric overlap analysis of same comparison in (E). Genes are ranked along axes by log2FC in indicated infection analysis (X: chronic HCV versus uninfected, Y: 18 m.p.i. chronic NrHV infection versus mock). Color indicates -log_10_ p-value for ranked list overlaps, with warmer colors indicating additionally significant ranked gene list overlap. Minimal permutation adjusted p-values are reported for each “directional” quadrant (*HCV down/ NrHV down* = 9 × 10^-114^, *HCV down/NrHV up* = 1, *HCV up/ NrHV down* = 0.92; HCV *up/ NrHV up* ≈ 0). (**G**) GSEA (HALLMARK gene sets) of significantly overlapping (HCV and NrHV) from (F, HCV *up/ NrHV up* and *HCV down/ NrHV down*). GeneRatio indicates number of genes in a given set divided by the total number of overlapping (input) genes.

To further assess the relevance of the NrHV model to human disease, we compared the gene expression changes elicited by chronic NrHV infection (mouse) with RNA-Seq data from liver biopsies of patients with chronic HCV infection versus uninfected individuals (*24, 25*). First, we used the HCV RNA-Seq data to derive two gene sets: “HCV_UP” and “HCV_DN”, representing genes up or downregulated during chronic HCV infection (versus uninfected controls) and mapped their mouse gene orthologues (**table S4**). Gene Set Variation Analysis (GSVA) indicated that NrHV infection resulted in significant enrichment of the HCV_UP set at all timepoints post-infection, and significant decreases of the HCV_DOWN set at two of three timepoints (**Fig. 3D**). We then focused on comparing HCV and NrHV 18 m.p.i. We performed a simple comparison of fold-change values for all expressed human-mouse orthologue (one-to-one) genes in HCV-infected vs uninfected patients and NrHV-infected vs mock-infected mice at 18 m.p.i. We observed a modest yet significant correlation (R = 0.43, p <2.2 × 10^-16^) of fold-change values (**Fig. 3E**). To account for differences in study design, sample size, and potential variability between infection datasets, we next compared the NrHV and HCV infection datasets by performing a rank-rank hypergeometric overlap analysis (*26*). Gene expression changes in NrHV and HCV exhibited highly significant similarities, particularly for genes increased in both NrHV and HCV infection (p.adj ≈ 0), but also for genes decreased in both infections (p.adj = 9 × 10^-114^). Discordant gene expression changes were not statistically significant (**Fig. 3F**). Gene sets associated with immune regulation (interferon alpha and gamma responses, allograft rejection, IL6-JAK-STAT3 signaling) were significantly enriched among genes upregulated by both infections, while gene sets associated with liver functions (bile acid metabolism, fatty acid metabolism, adipogenesis, xenobiotic metabolism) were significantly enriched among genes downregulated in both infections (**Fig. 3G**). These gene set patterns were confirmed when HCV and NrHV were analyzed independently (**fig. S6**).

Interestingly, although frank cirrhosis was not observed in NrHV-infected mice, we found a significant enrichment of hepatic stellate cell (HSC) activation signature derived from HCV patients with cirrhosis (*27*) which peaked at 18 m.p.i. in our model. This signature was previously reported associated with fibrosis severity and poor clinical outcomes in both cirrhosis and postresection HCC. These findings suggest that chronic NrHV infection progressively engages fibrogenic HSC programs reminiscent of human HCV pathology, supporting the utility of the model for studying fibrosis-driven disease progression (**fig. S7**).

Taken together, these results indicate that despite the cellular heterogeneity of the liver samples profiled and their disparate hosts, chronic human HCV and mouse NrHV infections exhibit gene expression signatures consistent with highly convergent biological processes, particularly those associated with immune functions and liver metabolism.

## Chronic NrHV infection causes spontaneous development of well-differentiated HCC in the context of chronic inflammation and advanced fibrosis

Chronic HCV infection results in significantly increased risk of HCC, which develops in the context of chronic hepatitis and advanced fibrosis/cirrhosis (*28, 29*). No infected mouse models of HCV-HCC are available and the artificial overexpression of the core protein - the only HCV protein shown to be oncogenic in mice - yields low (25%) HCC incidence (*30*), limiting its utility for mechanistic studies. In our model, 67% of NrHV-infected animals developed HCC in comparison to 4% of the mock-infected group, as confirmed by histological evaluation (**Fig. 4A**). Although variable across individual mice, plasma levels of alpha-fetoprotein (AFP) were significantly higher in infected mice with HCC (NrHV+HCC vs mock: p<0.0001; NrHV+HCC vs NrHV–HCC: p=0.0052) (**Fig. 4B**). There was a significantly higher incidence of HCC in NrHV-infected males than in infected females (p=0.01), mirroring the sex-associated differences observed in human HCC and chronic HCV infection (**Fig. 4C**). Combining two independent cohorts, 88% (15/17) of infected males and 43% (7/16) of infected females developed HCC at 18 m.p.i., while 6% of mock-infected males (1/17) and no mock-infected females developed HCC (0/8) (**Fig. 4C**). HCC was first observed 12 m.p.i. in 12.5% (1/8) of NrHV-infected males, and in no NrHV-infected females or mock controls at this time point (**fig. S8)**. In addition to higher HCC incidence, HCC tumors in males tended to be larger than in females (**Fig. 4D**). Mice that spontaneously resolved NrHV infection across the time course also developed HCC but at lower frequency than chronically infected mice (**fig. S9**).

**Fig. 4.**
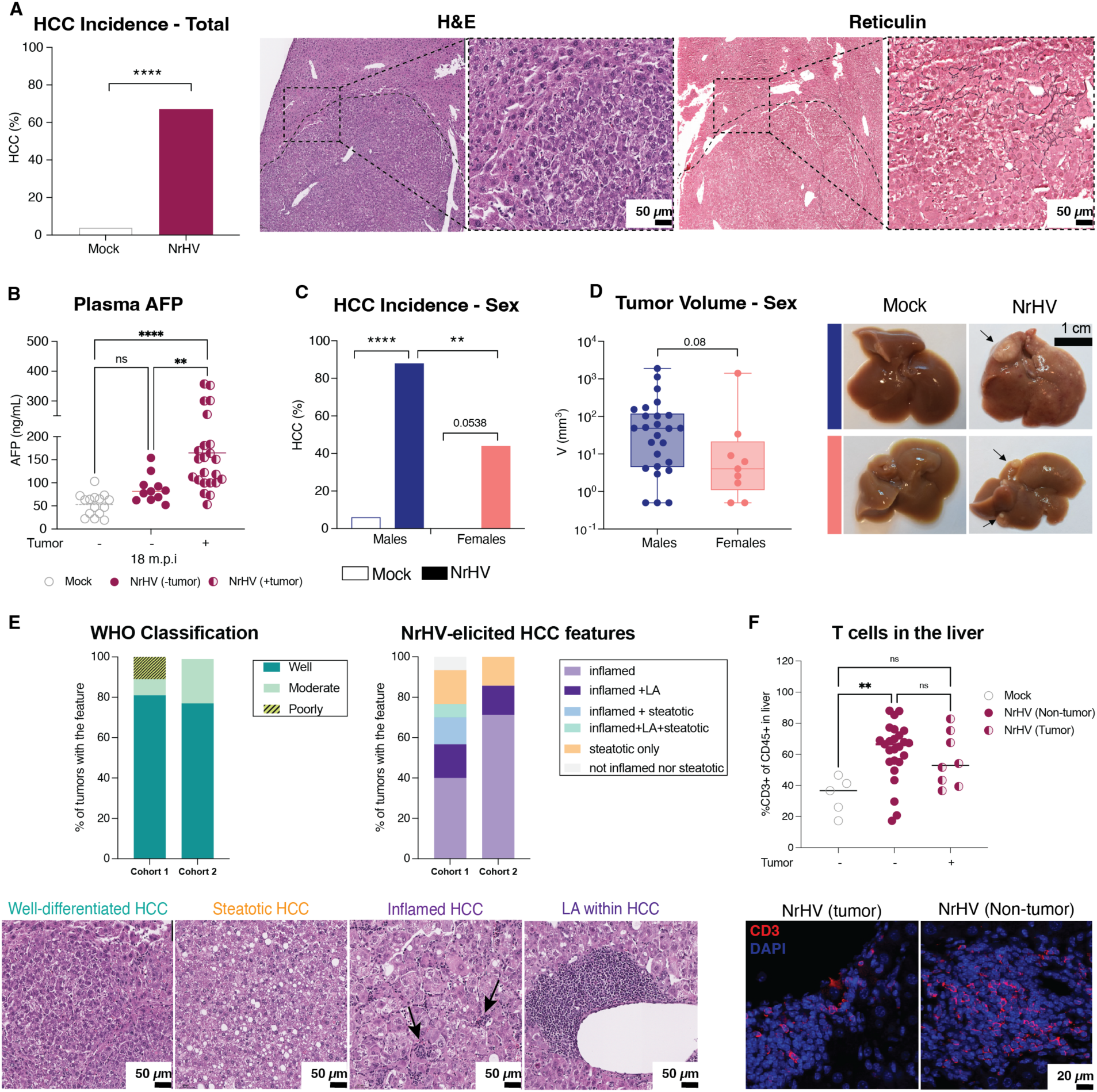
Chronic NrHV infection results in HCC which resemble histological features of “Non-Proliferation” class of human HCC. **(A)** HCC incidence in NrHV-infected mice 18 months post infection (m.p.i) in two independent experiments combined (total mice= 58; mock mice =25, NrHV mice=33). Statistics: Fisher’s exact test Mock vs NrHV ****p<0.0001. **(B)** Plasma AFP in NrHV-infected (Maroon) with HCC (half-solid); or not (solid) and mock-infected mice. Statistics: One-Way ANOVA Mock vs NrHV (+tumor) **** p<0.0001, NrHV(-tumor) vs NrHV(+tumor) **p=0.0052, ns: not-significant (p=0.38). **(C)** HCC frequency in male vs female mice. Dark blue: males, salmon: females, solid bars: infected, empty bars: mock. Statistics: Fisher’s exact test mock males (n=17) vs NrHV males (n=17) ****p<0.0001, mock females (n=8) vs NrHV females (n=16) p=0.0538 (ns); NrHV males vs NrHV females **p=0.01. **(D)** Tumor size and tumor gross view in NrHV-infected male vs female mice 18 m.p.i. Each dot represents one tumor, tumors found in the same mouse were counted separately (Total number of tumors = 57). Statistics: Fisher’s exact test *p=0.08, log transformation was applied to tumor volume for analysis. **(E)** Features of NrHV-elicited tumors found 18 m.p.i in two independent experiments. Main features are shown by H&E stain. **(F)** Percentage of T cells in the total leukocytes infiltrated in the livers of mock, NrHV-infected (non-tumor tissue) and NrHV-infected (tumor tissue) 18 m.p.i as defined by flow cytometry (top panel). Immunostaining showing CD3+ cells infiltrated in NrHV-elicited tumor (core) and adjacent liver tissue (bottom panel).

Histological assessments of NrHV-associated HCC from both cohorts revealed that more than 95% were well-differentiated, with trabecular patterns and preservation of hepatocyte morphology. Inflammatory and steatotic features were also commonly found with approximately 80% of tumors exhibiting an inflamed phenotype (high immune cell infiltration) and 25% exhibiting steatosis (**Fig. 4E**). Within these classifications, tumors were diverse, with highly variable degrees of differentiation and steatosis. Lymphoid aggregates (LA) were observed in a subset of tumors (25%) but were more frequently observed in adjacent non-tumor tissue (90% at 18 m.p.i.). Tumor and adjacent liver tissues from NrHV-infected mice were highly infiltrated by T cells in comparison to mock controls (**Fig. 4F**). Human HCC are highly heterogeneous, with histopathological characteristics often associated with specific etiologies. Our results show that NrHV HCC histologically mirrors the ‘non-proliferation’ class of human HCC, which is associated with HCV infection as well as metabolic-associated steatohepatitis (MASH), and chronic alcohol use(*2*).

## NrHV-elicited HCC is molecularly heterogenous but frequently exhibits convergent mechanisms in well-differentiated HCC

To characterize the molecular heterogeneity in NrHV-elicited HCC, we performed RNA-Seq on tumors and matched adjacent liver tissue collected at 18 m.p.i. from both cohorts. Despite highly variable transcriptional profiles across tumors (**fig. S10A**), differential expression analysis identified 261 upregulated and 217 downregulated genes in tumor vs. adjacent tissue (**fig. S10B**). Gene set enrichment (*23*) **(fig. S10C)** indicated consistent increased expression of genes associated with cell cycle, proliferation and immune signaling pathways in tumor, and decreased expression of genes associated with metabolic pathways (including bile acid metabolism and oxidative phosphorylation) in some of the tumors evaluated (**Fig. 5A, panel 1**). Of note, several pathways found to be increased in tumors were also elevated in the adjacent non-tumor tissue (**Fig. 3C** above), consistent with the ‘field effect’ theory. This concept proposes that chronic injury or infection induces widespread molecular alterations in the liver, creating a microenvironment conducive to tumor development(*31, 32*). While some pathways were consistently enriched across tumors, others varied across samples, and in some cases, across different tumor samples from the same animal, similar to the inter and intra-heterogeneity commonly observed in HCC patients. This intra-host heterogeneity again supports the idea that tumors can arise independently in a cancer-prone microenvironment as previously suggested in subsequent *de novo* tumors in human cirrhotic livers (*31, 33*). Cophenetic correlation, which quantifies how strongly ssGSEA values contribute to clustering structure, revealed metabolism-related pathways (e.g. fatty acid, xenobiotic metabolism, and adipogenesis) and cytokine signatures (IL6 JAK STAT3 signaling and IFN response) as key drivers of tumor sample heterogeneity distinguishing two broadly distinct groups of NrHV-elicited tumors (**Fig. 5A, panel 2**).

**Fig. 5.**
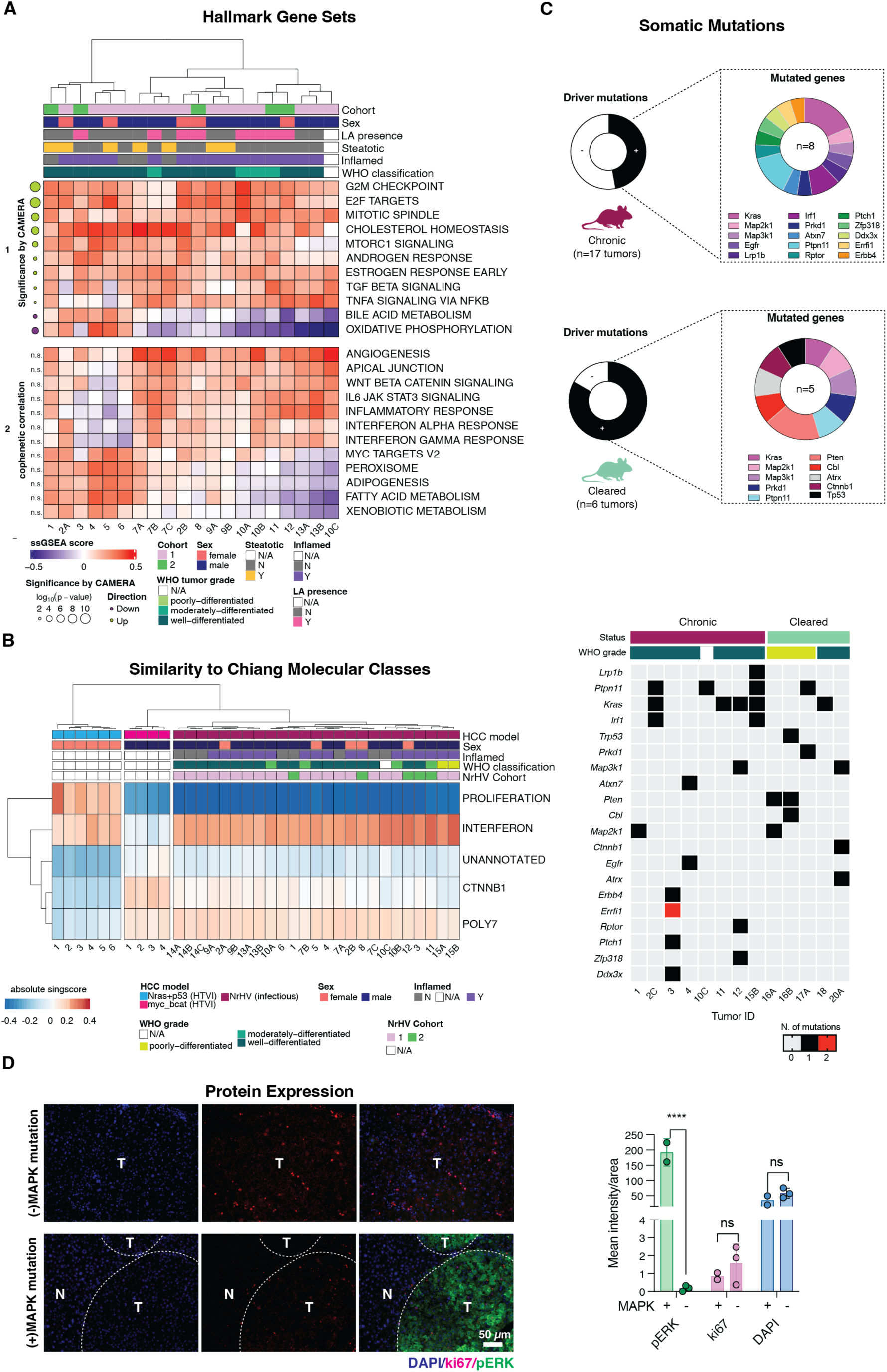
Molecular analysis of NrHV-elicited tumors reveals heterogeneity and similarity with human “Interferon” molecular subclass. **(A)** Panel 1: Heatmap depicting ssGSEA scores subset on significantly enriched Hallmark gene sets shared across tumor vs. adjacent tissue in both cohorts. The dot plot shows the corresponding significance and direction of enrichment for each gene set. Color indicates enrichment (light green) or de-enrichment (purple) of gene sets in tumor vs adjacent tissue and dot size indicates significance. Panel 2 depicts gene sets not significant in Panel 1 that contribute most to the clustering structure of the full Hallmark gene set ssGSEA matrix (**Fig S11C**). All tumors from infected mice with paired adjacent liver tissue passing RNA-Seq filtering criteria were included (n=21 tumors, from 13 mice) **(B)** Similarity Scores (singscore) of NrHV-elicited tumors compared to the Chiang Human HCC molecular subclasses. All tumors from infected mice with high quality RNA were included in this analysis irrespective of adjacent pairing (n=25 tumors, from 15 mice). HTVI samples from ≈-catenin and p53 mutant tumor models were used as controls for the “proliferative” and “≈-catenin” molecular classes. **(C)** Putative driver mutations found in individual tumors detected by targeted DNA-seq (mouse IMPACT). A total of 17 tumors were sequenced from chronically infected mice (chronic) and 6 from cleared mice (cleared). Donut charts indicate presence of somatic mutations in the total sequenced tumors in black (present), white (absent) (chronic: present in 8/17; cleared: present in 5/6 sequenced tumors) and in which genes they were present (each mutation is represented by a different color). Categorical heatmap (below) demonstrates the collection of mutations in each individual tumor. Tumors with no putative mutations are not indicated in the heatmap but are included in the “driver mutations” donut chart. **(D)** Immunohistochemistry from tumor and adjacent liver tissue showing the effect of putative mutations on Ki67 (proliferation, red) and pERK (MAPK activation, green) expression. Nuclei (DAPI) are shown in blue.

To further contextualize NrHV-elicited tumors, we classified their transcriptional profiles using Chiang HCC molecular definitions (*34*), which categorizes HCV-associated HCC into five molecular subclasses: “CTNNB1” (beta-catenin), “Proliferation”, “Interferon”, “Polysomy of Chromosome 7”, and an “Unannotated Subclass”. Using the singscore tool(*35*) applied to normalized RNA-Seq expression data from both cohorts, we compared NrHV tumors with public datasets from common mouse HCC models induced via hydrodynamic tail vein injection (HTVI) of p53^del^/Nras^G12V^-(representing poorly differentiated tumors with low T cell immune infiltrate)(*36*) and β-catenin/Myc overexpression (representing moderately-differentiated with moderate T cell infiltrate) mutated tumors (*37*). As expected, p53^del^/Nras^G12V^ tumors were classified in the “Proliferation” subclass while β-catenin/Myc tumors were assigned to “CTNNB1”. In contrast, NrHV tumors were more similar to the ‘Interferon’ subclass, regardless of histological inflammation status, likely reflecting persistent antiviral signaling in the setting of chronic viral infection **(Fig. 5B**). Scores were more variable for NrHV than for the transgenic models, likely reflecting stochastic events associated with the natural development of HCC in our model and the molecular heterogeneity previously described. We also expanded our analysis to other two major human HCC classifications (Hoshida and Boyault)(*31, 38*) with similar results (**fig. S11**). Tumors from cleared mice, although still generally similar to HCC from chronic mice, formed separate clusters (**fig. S12**). Importantly, the gene set clusters and variation within could reflect differences in intrinsic tumor biology, immune cell infiltrates, and/or other differences in the relative frequency of component cell types in bulk liver tissue specimens.

## NrHV-elicited HCC reflects genetic heterogeneity of human HCC

To identify genetic drivers of NrHV-induced HCC, we performed targeted DNA sequencing of 23 independent tumors (17 from chronically infected mice and 6 from cleared mice) using the Mouse MSK-IMPACT panel. Mouse MSK-IMPACT is a curated version of the clinically validated MSK-IMPACT assay adapted for mouse models, covering the coding regions of approximately 600 cancer-associated genes (**table S5**). As in humans, NrHV-HCC exhibited a highly heterogenous genetic profile. Interestingly, while observed at low frequency, mutations in genes commonly associated with HCV-HCC such as *Ctnnb1* (≈-catenin), *p53*, and *Pten* were identified(*39*). *Pten* and *p53* mutated tumors were poorly differentiated, while the single tumor with *Ctnnb1* mutation was well-differentiated, consistent with associated features in human HCC. In addition, several tumors harbored mutations affecting the RAS/MAPK pathway, including *Kras*^G12V^, *Kras*^Q61R/H^, *Ptpn11*^E69K^, *Map2k1*^C121S^, *Map3k1*^F349L^ and homozygous loss of *Erffi* (**Fig. 5C**). Supporting the functional relevance of these mutations, tumors with RAS/MAPK alterations displayed elevated levels of phosphorylated ERK (pERK), consistent with pathway activation (**Fig. 5D**). Although *Kras* mutations are rare in human HCC patients, the RAS/MAPK pathway is overactivated in more than 20% of cases (*40*). Notably, tyrosine kinases emerge as key players in this model, aligning with the clinical relevance of tyrosine kinase inhibitors (TKIs), which remain a cornerstone of standard chemotherapy for HCC in humans(*41*). The mutational landscape of NrHV-induced tumors further underscores the heterogeneous nature of hepatocellular carcinoma (HCC), while also revealing strongly convergent pathways which can be now explored to define novel tumor vulnerabilities.

## Conclusion

Overall, we demonstrate that the extended duration of chronic NrHV infection in immunocompetent B6 mice recapitulates key features of chronic HCV infection in humans, including sustained hepatitis and progressive liver fibrosis. Remarkably, chronic NrHV infection leads to spontaneous HCC development without additional experimental intervention such as chemical insult or genetic manipulation. As such, this immunocompetent mouse model addresses many of the significant limitations of prior HCC and HCV animal systems and thereby establishes a novel research tool for liver pathogenesis studies. Importantly, transcriptional alterations were observed not only in tumors but also in adjacent non-tumor liver tissue, consistent with a field effect that may predispose the liver to malignant transformation. This highlights the utility of the model for studying early molecular changes that precede tumor development. Future studies can take full advantage of many well-established genetically engineered B6 models enabling for detailed mechanistic investigations of hepacivirus pathogenesis, immune mediated inflammation, fibrosis and tumor initiation. Further exploitation of this model may disentangle the relative contributions of viral infection and immune dysfunction to HCC oncogenesis. Finally, with its strong parallels to chronic HCV, the chronic NrHV system provides an invaluable and physiologically relevant platform for preclinical testing of prophylactic strategies and therapeutic interventions targeting virus-associated HCC.

## Materials and Methods

### Mouse Infection Studies

All mice used in this study were housed and bred at The Rockefeller University Comparative Biosciences Center in accordance with the NIH Guide for the Care and Use of Laboratory Animals and approved by The Rockefeller University Animal Care and Use Committee. All animals included in this study were CD4 T cell depleted as previously described (*8*). In cohort 1 (n=75) 10- to 12-week-old wild-type C57BL6/J (B6) mice were grouped in 4 cohorts as follows: NrHV-infected females (n=30), NrHV-infected males (n=30), mock-infected females (n=5); mock-infected males (n=10). In cohort 2 (n=96), B6 mice underwent the same procedures as described above. Age and gender-matched animals were injected with 100 µL of DPBS and used as mock-infected controls. One day prior to experiment launch, mice were weighed and bled for baseline references. For post infection analysis, mice were bled monthly. Monthly blood analyses included: NrHV viremia and alanine transaminase. Infected mice with undetectable viremia at month 1 were excluded from all analysis as they were considered unsuccessful launch (refer to table S1 and S2). Alpha-feto protein levels were measured in blood samples collected at 18 m.p.i. General mouse activity was monitored weekly. Subgroups of infected and mock-infected animals were euthanized by CO_2_ 18 months post-infection in cohort 1. In cohort 2, of 58 infected mice, 15 mice (7 females/8 males) were euthanized 7 m.p.i; 19 mice (10 females/9 males) were euthanized 12 m.p.i and 14 mice (8 females/6 males) were euthanized 18 m.p.i. For each time point, organs from at least 5 mock controls of each sex were also collected. For organ harvest, animals were euthanized and immediately perfused with DPBS (Gibco) by cardiac perfusion for 3 minutes using a peristaltic pump to remove blood from the liver. Livers were resected, weighed and slices were either stored in RNAlater (Thermo), fixed in 10% buffered formalin, 4% paraformaldehyde (PFA) or flash frozen with liquid nitrogen and stored at −80C until subsequent analysis. Mice that died prior to 18 m.p.i. endpoint (cohort 1) were used to determine survival probability. For both cohorts, mice were humanely sacrificed no later than 18 m.p.i. Mice that spontaneously cleared NrHV infection were excluded from analyses, unless otherwise specified. Experimental design for cohort 1 and cohort 2 is depicted in Figure 1A and 2A, respectively. Humane endpoints included: tumors exceeding 2000mm^3^, inactivity, weight loss higher than 10% and rectal prolapse.

### Recombinant Virus Production

DNA from the infectious clone NrHV-B_SLIS_, carrying the T190S, V353L, F369I and N550S mouse adaptive mutations(*11*) was linearized using MluI and purified using DNAclean and concentrator (Zymo). RNA was transcribed from 1 μg of MluI-linearized DNA using T7 RiboMAX Express Large Scale RNA Production System (Promega). Template DNA was degraded using RQ1 DNAse for 30 min at 4°C. Subsequently, RNA was purified using RNAeasy mini kit (Qiagen). Five 4-week-old NRG mice were injected intra-hepatically using 10 μg of RNA in a maximum volume of 50 μL of RNA diluted in PBS. A terminal bleeding was performed one week post infection and serum from all mice was pooled, titrated by RT-qPCR, sequenced and stored in −80°C freezer (passage #0). To amplify the stock, new cohorts of seven 4-week-old NRG mice were infected intravenously with 10^4^ genomic equivalents (GE) of passage #0 per mouse. The same harvest protocol was followed and serum was titrated, sequenced and stored at −80°C (passage #1). All experiments were performed using virus passage #1. Sequencing primers, RT-qPCR primers and probe for NS3 standard curve were as described(*13*).

### Blood collection and processing

Blood was collected in EDTA tubes (BD) monthly and at terminal harvests. Tubes were centrifuged 9,600 x g, at room temperature for 5 minutes, plasma transferred was to a new tube and stored at −80°C for further analysis.

### RNA isolation from serum and qRT-PCR

RNA was isolated from 10 µL of mouse plasma using High Pure Viral Nucleic Acid kit (Roche) following the manufacturer’s instructions. To quantify viral genomic RNA, reverse transcription – quantitative polymerase chain reaction (qRT-PCR) was performed using 2 µL of eluted RNA, TaqMan Fast Virus 1-step Master Mix (Applied Biosystems) and primer/probes targeting NS3 (IDT). Reactions were performed in QuantStudio^TM^ 3 (Thermo). The following RT-PCR program was run: 50°C for 30 minutes, 95°C for 5 minutes, followed by 40 cycles of 95°C for 15 seconds, 56°C for 30 seconds, and 60°C for 45 seconds, then 40°C for 10 seconds. The sequence of the NrHV NS3 primers and probes were: Forward: 5’-GTCCGACCGAACATCCACTGT −3’, Reverse: 5’-GCATCCAGCAGGTTCTTCGC −3’, Probe: [6-FAM]CTCACGTACATGACGTACGGCATG[BHQ1a-6FAM]. NrHV standard curve RNA was produced as previously described by (*13*).

### ALT analysis

Longitudinal analyses of ALT activity levels in the plasma were carried out using the Alanine Transaminase Assay kit (Abcam) according to the manufacturer’s instructions. ALT activity was measured 1, 6, 12- and 18-months post infection (m.p.i).

### AFP analysis

AFP levels in the plasma were measured with the Mouse alpha-fetoprotein/AFP Quantikine ELISA Kit (Bio-Techne). Plasma samples were diluted 20x in calibrator diluent. AFP was measured for all NrHV and mock-infected mice harvested 18 m.p.i in cohort 1 and 2.

### Histology

To determine the pathologic effects of NrHV infection in the liver over time, organs collected at 7-, 12- and 18- m.p.i were prepared using standard histopathology techniques. A representative fraction of the median right and left lobe of the liver was formalin fixed (3-4 days) transferred to 70% ethanol and processed using Leica Peloris with routine dehydration, clearing and infiltration schedule (50-minute steps). Tissues were paraffin embedded and sectioned at 4 micron thickness. H&E staining was performed on a Leica ST5020 automated stainer and scanned with the Leica SCN 400F. To assess liver fibrosis, livers were stained manually using Picro Sirius Red (Abcam). For confirmatory pathology of HCC, reticulum stain was performed using HT102 (Sigma-Aldrich), following the manufacturer instructions. Immunohistochemical (IHC) or immunofluorescent staining of selected liver tissue sections were performed for alpha-smooth muscle actin, cleaved caspase-3, Ki67, cluster differentiation 3 (CD3) and phosphorylated ERK. Antibodies catalogs and concentrations: Alpha-smooth muscle actin (α-SMA) (Abcam #ab32575, 1:250), Caspase-3 (Cell Signaling Technology #9661, 1:250), ki67 (Cell Signaling Technology #12202, 1:500), CD3 (Abcam #ab5690; 1:200, anti-rabbit AF647 – Thermo # A-21244, 1:500); pERK (Cell signaling Technology #4370; 1:400). Tissue sections were evaluated by a Board-Certified human pathologist (*Pathology and Anatomic Pathology* by the American Board of Pathology) and scores for fibrosis and hepatitis in mice were developed based on criteria of METAVIR score for inflammation and ISHAK score for fibrosis as described in **table S3** (*22*). Liver samples from our two independent cohorts classified by gross view as tumors by initial researcher inspection were submitted to histology (H&E/ reticulum/PSR) and then evaluated by two independent pathologists. A designation of “HCC” was assigned if diagnosed by at least one pathologist (refer to **table S2** for details). The number of tumors per mouse was recorded after pathological confirmation. Tumor volume was determined by caliper measurements at harvest and included only if HCC was confirmed by histology. Cleared mice were defined as those with undetectable viremia and virus RNA in the liver at the time of harvest. Mice initially included in the infected group but with no detectable viremia within one-month post-infection were considered a failed launch and were excluded from the analysis. Staining quantifications were performed using ImageJ 1.54g, Java 1.8.0_345 (64-bit).

### RNA/DNA isolation from liver

RNA and DNA were isolated from nitrogen pulverized liver slices which were previously flash-frozen in liquid nitrogen. Isolation was performed using the Allprep Qiagen kit (QIAGEN) following manufacturer’s instructions, with the addition of DNAseI to the original RNA isolation protocol. Some samples presented with suboptimal A260/230 ratio, therefore all samples were purified using either RNA mini kit (Zymo Research) or DNA purification kit (Zymo Research). Samples were quantified by Qubit and assessed for quality by Agilent BioAnalyzer. RNA samples with RNA Integrity Number (RIN) above 7 were included in downstream library preparation and sequencing workflows.

### Immune cell isolation from liver, immunostaining, and flow cytometry

Dissociated immune cell suspensions were obtained from livers (tumor and adjacent tissue) as previously described(*42*). After processing, cells were counted by hemocytometer and 2-3 million cells were labeled for flow cytometry analysis. Extracellular labeling was performed in FACS buffer (PBS + 2%FBS + 2mM EDTA + 0.05% sodium azide) for 30 min at 4C using anti-CD45 BV510 (clone: 30-F111 Biolegend −1:200) and anti-CD3-APC-Fire810 (clone: 17A2 Biolegend - 1:200) followed by fixable Live Dead Blue (Thermo-Fisher – 1:1000) staining in PBS+1%EDTA for 15 min. Cells were fixed using 2% PFA for 20 minutes, washed in FACS buffer. Samples were acquired the following day on the Aurora Spectral Flow Cytometer (Cytek Biosciences) and data were analyzed in Flowjo v10.

### RNA-Seq analysis of NrHV- and mock-infected mouse livers

mRNA sequencing (RNA-Seq, polyA selection, strand-specific) was performed by Novogene with the NEBNext Ultra II RNA Library prep kit (New England Biolabs). RNA-Seq libraries were sequenced on the Illumina Novaseq platform, with paired end configuration (150nt x 2), with a targeted read depth of 20 million read pairs per library. Sequence quality was assessed with FastQC (v0.11.9) and adaptor sequences were trimmed with fastp (v.0.23.2)(*43*). Read mapping was performed with Salmon (v.1.4.0)(*44*) in selective alignment mode(*45*). Reads were mapped to mouse reference transcriptome (GRCm39, release M32), appended with NrHV clone B_SLIS_ sequence(*11*)(GenBank accession ON758386.1). Subsequent analyses were conducted within the R statistical framework (v4.0.3). Differential gene expression analysis was performed on Salmon-estimated counts with edgeR (v.3.32.1)(*46*), and gene set enrichment testing was performed with CAMERA(*47*). For visualization and clustering purposes, gene expression count values were normalized and variance-stabilized with the vst function in the DESeq2(*48*) package (v.1.30.1), and Gene Set Variation Analysis (GSVA) and single sample Gene Set Enrichment Analysis (ssGSEA) scores were calculated using the GSVA (*49*) package (v.1.38.2). Significance comparisons of GSVA values were performed as described below (‘Statistics’), using the limma package (v3.46.0) and sva package (v3.38.0). HCV infection gene sets were constructed by conducting differential gene expression analysis (edgeR) on infected vs. control liver datasets (Database Accessions : E-MTAB-6863(*25*) (ArrayExpress), GSE126848(*50*) (GEO), GSE135251(*47*) (GEO), GSE142530(*51*) (GEO), GSE84346(*24*) (GEO), SRP170244(*52*) (SRA)), including the top 200 upregulated (HCV_UP) and top 200 downregulated (HCV_DOWN) ranked by adjusted p value. Cophenetic correlation was calculated on the clustering hierarchy of the ssGSEA matrix for all 50 Hallmark gene sets using the stats (v4.0.3) package. Rank-rank hypergeometric overlap (RRHO) and associated statistical gene-gene correlation analyses was enabled by the RedRibbon package (v1.3.1)(*26*). Subsequent gene set enrichment analysis on significantly overlapping genes was performed using gprofiler2 (v0.2.3)(*53*). Raw data and processed data are available at the NCBI GEO repository under accession GSE265927.

### Targeted DNAseq

Memorial Sloan Kettering-Integrated Mutation Profiling of Actionable Cancer Targets (MSK-IMPACT) is a custom hybridization capture–based assay encompassing all genes that are druggable by approved therapies or are targets of experimental therapies being investigated in clinical trials at Memorial Sloan Kettering Cancer Center (MSKCC), as well as frequently mutated genes in human cancer (somatic and germline mutations). MSK-IMPACT is capable of detecting sequence mutations, small insertions and deletions, copy number alterations and select structural rearrangements. M-IMPACT_v2 (mouse version) assay covers more than 600 known cancer driver genes (**table S5**). For the detection of somatic genomic alterations in NrHV-elicited tumors, we extracted genomic DNA from tumor and normal uninfected liver samples and gDNA was enriched for targeted regions using a solution-phase hybridization-based exon capture as previously described(*54*). Captured DNA fragments were sequenced on an Illumina HiSeq 2500 as paired-end 100-bp reads with intended 500× coverage for all reads. Sequencing analysis was performed by the MSKCC Bioinformatics Core. In brief, processed FASTQ files were mapped to the mouse reference genome mm10 (GRCm38) using BWA-MEM. Local realignment and quality score recalibration was performed using Genome Analysis Toolkit (GATK) according to GATK best practices(*55*). Sample variant calling was performed on tumor samples and normal samples to identify point mutations/single nucleotide variants (SNVs) and small insertions/ deletion (indels). MuTect (version 1.1.7)(*56*) was used for SNV calling and SomaticIndelDetector, a tool in GATKv.2.3.9, was used for detecting indel events. Variants were subsequently annotated using Annovar, and annotations relative to the canonical transcript for each gene (derived from a list of known canonical transcripts obtained from the UCSC genome browser) were reported. Variant allele frequency (VAF) was determined by calculating the fraction of variant reads for a specific out of the total reads at that location. Raw data and processed data are available at the NCBI GEO repository under accession GSE265927.

### Statistics

Statistics were calculated using GraphPad Prism, SAS Studio, and R. Kaplan-Meier method and log-rank test were used to compare overall survival. Mann-Whitney test and Kruskal-Wallis with Dunn’s post-hoc test was used for comparing continuous variables. Fisher’s exact test was used to compare proportions of tumor development. Mixed-effects model for repeated measures was used to analyze tumor volume. One-way ANOVA with Turkey’s post-test was used to analyze AFP, spleen and liver weight 18 m.p.i. Two-way ANOVA with Sidák’s post-test was used to analyze viremia and ALT levels over time in males vs females. Significance comparisons of GSVA values were performed using the recommended linear-model–based testing approach (with empirical-Bayes shrinkage) in R to compare overall gene set enrichment in chronic vs. naive samples at each timepoint. For all analysis, p-values < 0.05 were considered statistically significant.

## Supplementary Figure and Tables

**Fig. S1.**
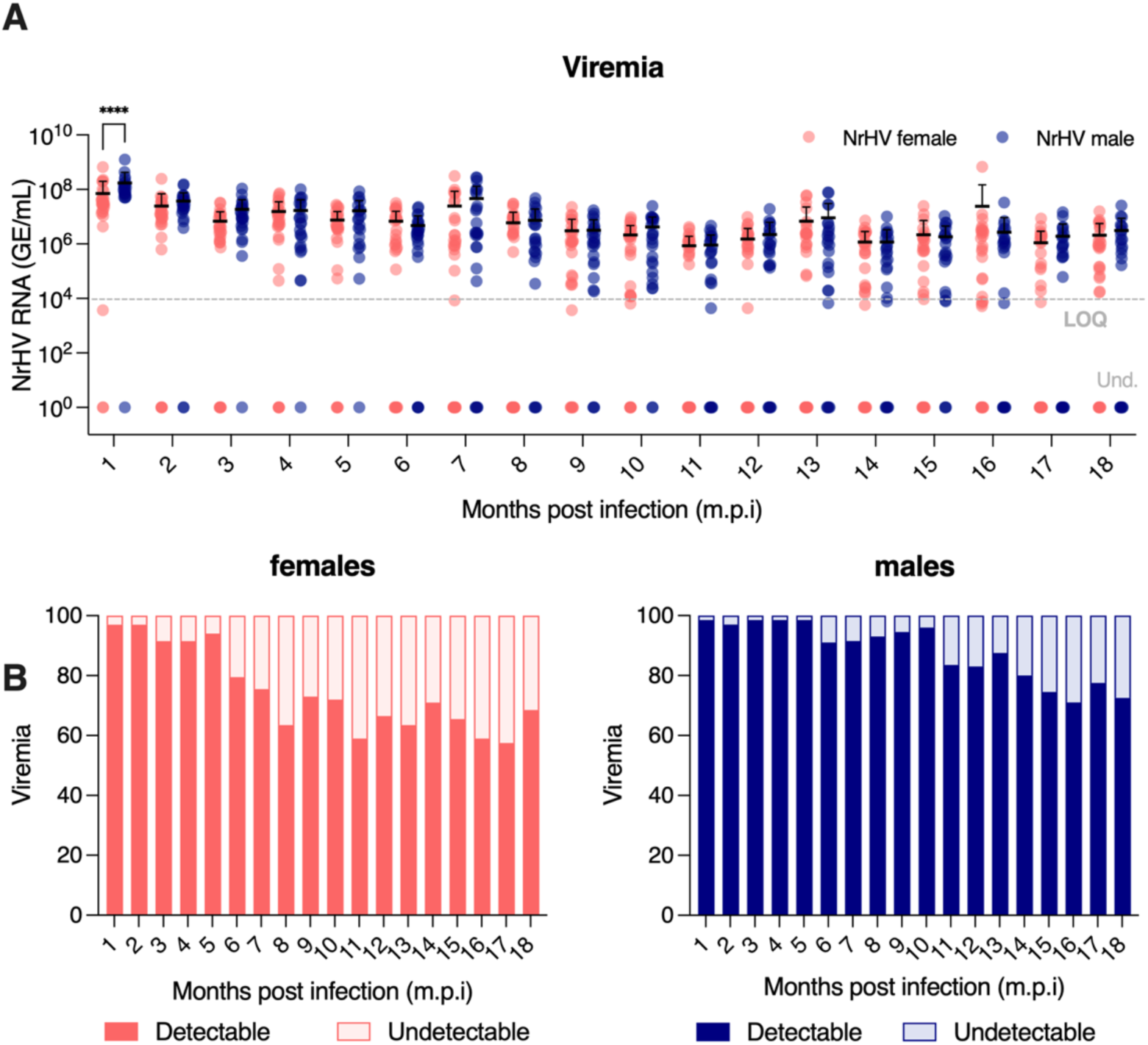
NrHV viremia comparison between male and female mice. **(A)** Viremia (mean ± SD) in NrHV-infected animals during 18 months of experimentation. Statistics: Two-way ANOVA: ****p<0.0001 males vs females. Differences in viremia between males and females after 1-month post-infection (m.p.i) were not significant. Mice which survived the 18 months of experimentation from cohorts 1 and 2 were included in the analysis (n=30 females, 26 males). **(B)** Percentage of mice with detectable (dark color) vs undetectable (light color) viremia by qPCR per timepoint. Bars indicate the percentage of total assessed animals. The initial number of infected animals was the same (n=58 per sex), and mice that died throughout the experiments were excluded from the analysis. Salmon: females, blue: males.

**Fig. S2.**
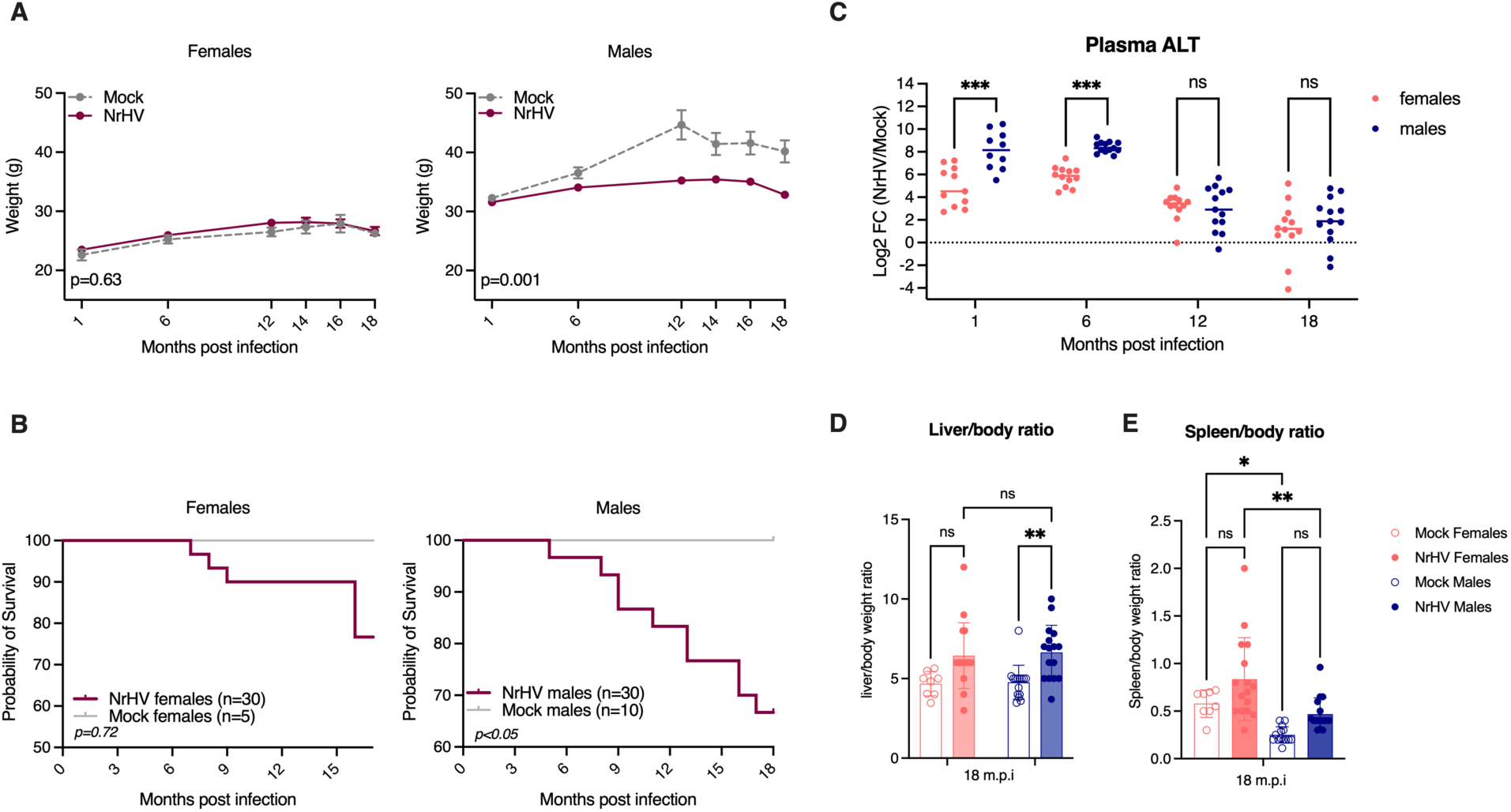
General features comparing males and females. **(A)** Body weight (mean ± SEM) of NrHV-infected vs mock controls over time. Statistics: Mixed effects model for repeated measures, Log transformation was applied prior to analysis, overall sex effect (p<0.0001). Female mice: Status p=0.63, Time p<0.0001, Status*Time p=0.81; Male mice: Status p<0.0001, Time p<0.0001, Status*Time p<0.0001 (n= 30 NrHV females, 5 mock females, 30 NrHV males, 10 mock males) **(B)** Kaplan-Meier survival analysis. Maroon lines: NrHV-infected, gray lines: mock controls. Statistics: Log-rank (Mantel-Cox) test p=0.72 for female comparison, p<0.05 for male comparison. **(C)** Log2 Fold change of alanine aminotransferase (ALT) activity of NrHV vs mock measured in plasma of mice over time. Statistics: Two-way ANOVA, ***p<0.001 male vs female (1 m.p.i n=11 NrHV females, 10 NrHV males; 6, 12 and 18 m.p.i n=12 of each sex). **(D)** Liver/body weight ratio (boxplot, min to max) of NrHV-infected females vs males 18 m.p.i. Statistics: One-way ANOVA, **p<0.01 NrHV male vs mock male, ns= not significant (n=12 NrHV females, 12 NrHV males, 3 mock females, 10 mock males). **(E)** Spleen/body weight ratio (boxplot, min to max) of NrHV-infected females vs males and age matched controls 18 m.p.i. Statistics: One-way ANOVA, **p=0.0015 NrHV female vs NrHV male, *p=0.04 mock male vs mock female, ns: not significant. All samples are from cohort 1. To increase statistical power organ/body weight ratios are calculated considering cohort 1 and 2 for mice alive 18 m.p.i (n=16 NrHV females, 17 NrHV males, 8 mock females, 16 mock males).

**Fig. S3.**
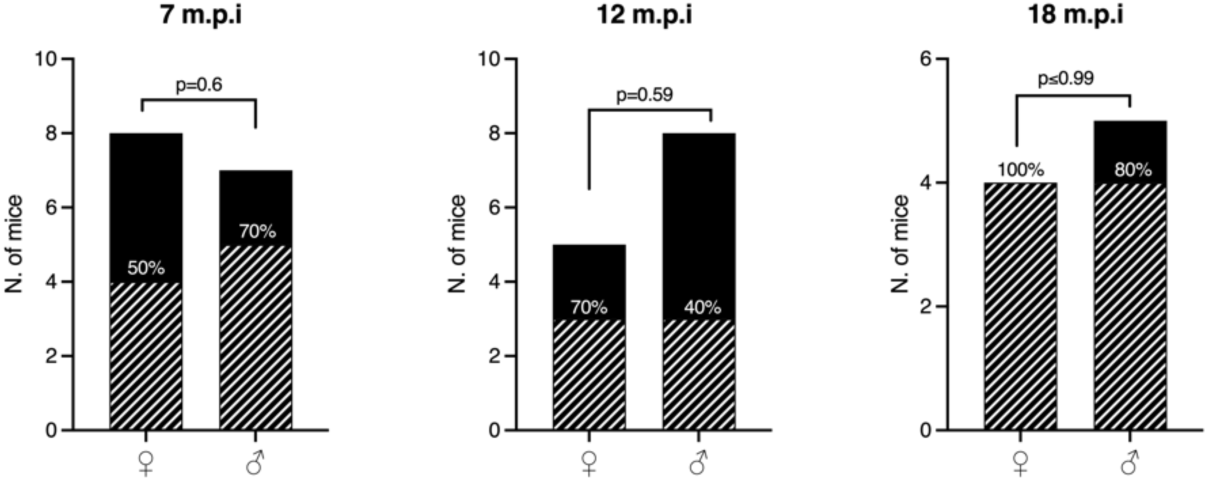
Percentage of NrHV-infected mice with lymphoid aggregates (LA) in the liver over time. Livers resected 7- (n=8 females, 7 males), 12- (n=5 females, 8 males), and 18- months post-infection (n=4 females, 5 males) were stained with H&E and analyzed blindly by a Board-Certified human pathologist (*Pathology and Anatomic Pathology* by the American Board of Pathology). Bars indicate number of mice with (dashed) and without LA (solid). Percentage indicates frequency of LA in each group. Statistics: Fisher’s exact test, all p>0.59 (not-significant) males vs females at all time points. All samples are from cohort 2.

**Fig. S4.**
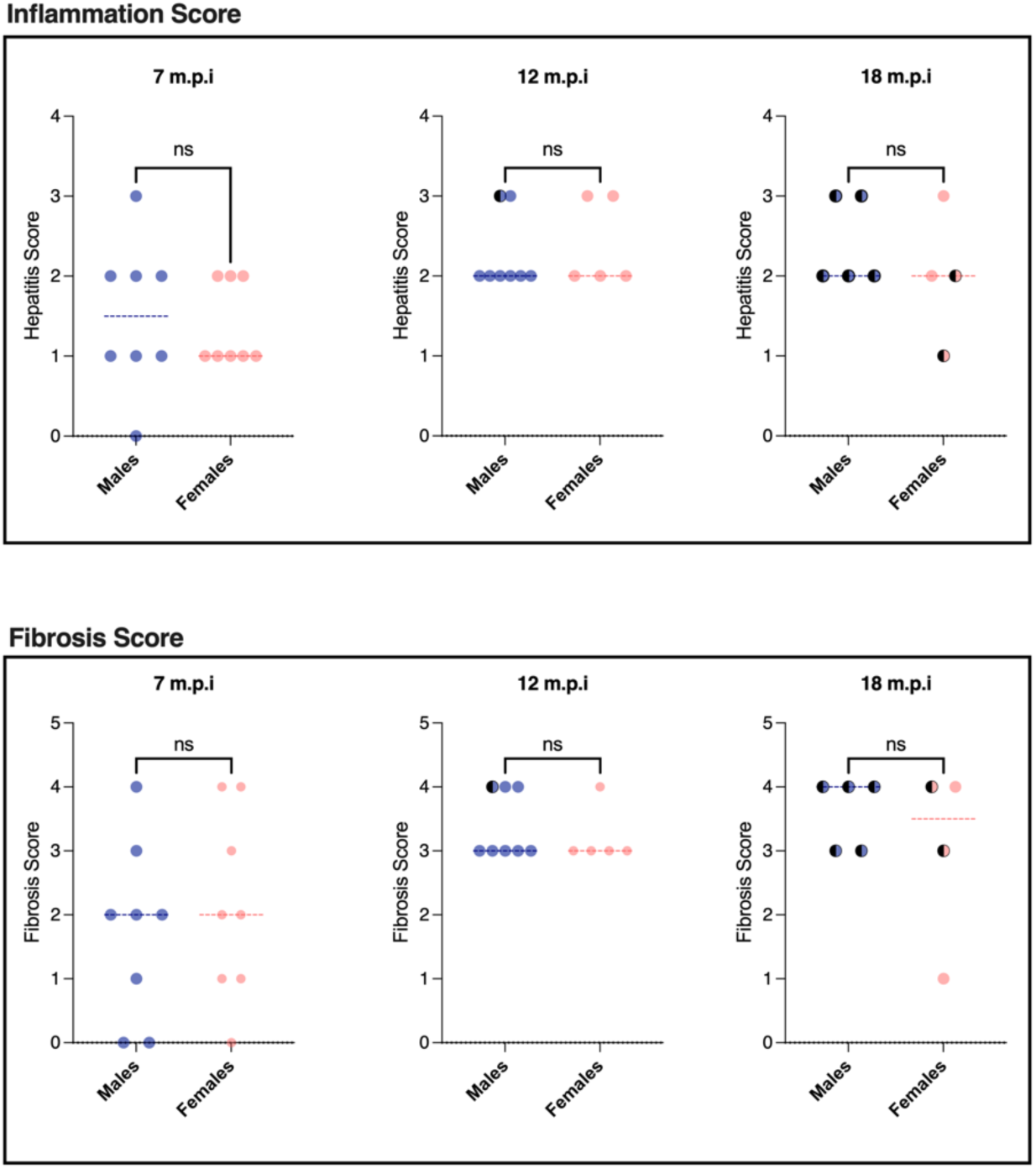
Fibrosis and hepatitis activity (inflammation) scores in NrHV males vs females. Top panel (Fibrosis). Bottom panel (Inflammation). Livers resected 7- (n=8 per sex), 12- (n=5 females, 8 males), and 18-months post-infection (n=4 females, 5 males) in Cohort 2 were submitted to H&E or PicroSirius Red (PSR) staining and blindly analyzed by a a board-certified human pathologist (*Pathology and Anatomic Pathology* by the American Board of Pathology). Scoring criteria are described in **table S3**. Statistics: Unpaired t test. Females: salmon, males: dark blue. Mice which developed tumors are represented as half-solid symbols. All samples are from cohort 2.

**Fig. S5.**
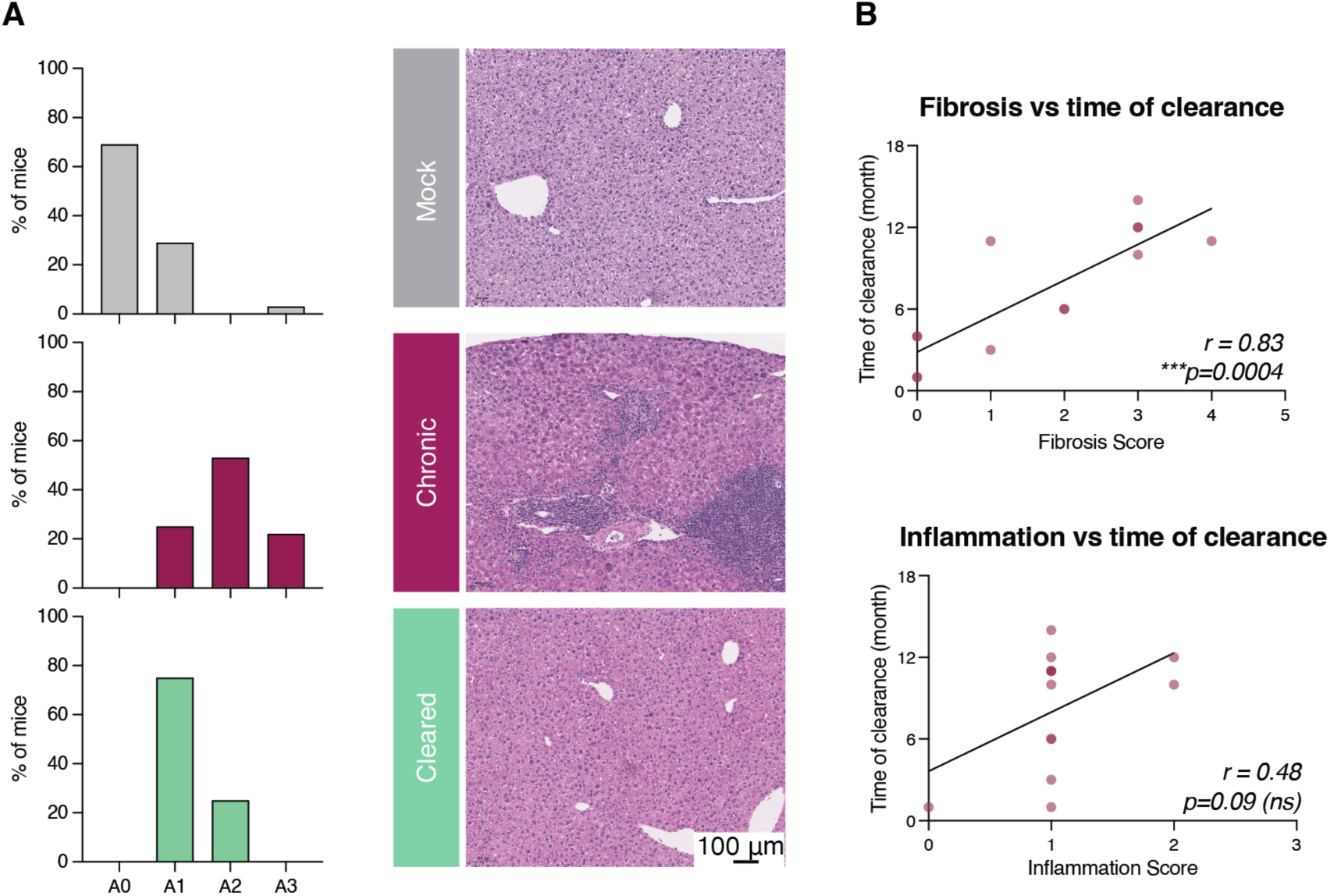
Inflammation and Fibrosis Scores upon clearance. **(A)** Inflammation Scores of livers comparing mock (gray, n=35); chronic NrHV (maroon, n=36) and resolved (green, n=13) infection. H&E staining in representative liver sections for each group. (**B**) Pearson correlation was used to define the linear relationship between time of spontaneous clearance and fibrosis and/or inflammation score. The analysis yielded a Pearson r= 0.83, p=0.0004*** (fibrosis score) and 0.48, p=0.09 (ns) (inflammation score) indicating that time of clearance has a strong correlation with fibrosis score but a moderate correlation with inflammation score. Analysis included all mice in cohort 2 that spontaneously cleared NrHV irrespective of time of clearance and across all harvest timepoints (n=13). All samples are from cohort 2.

**S6.**
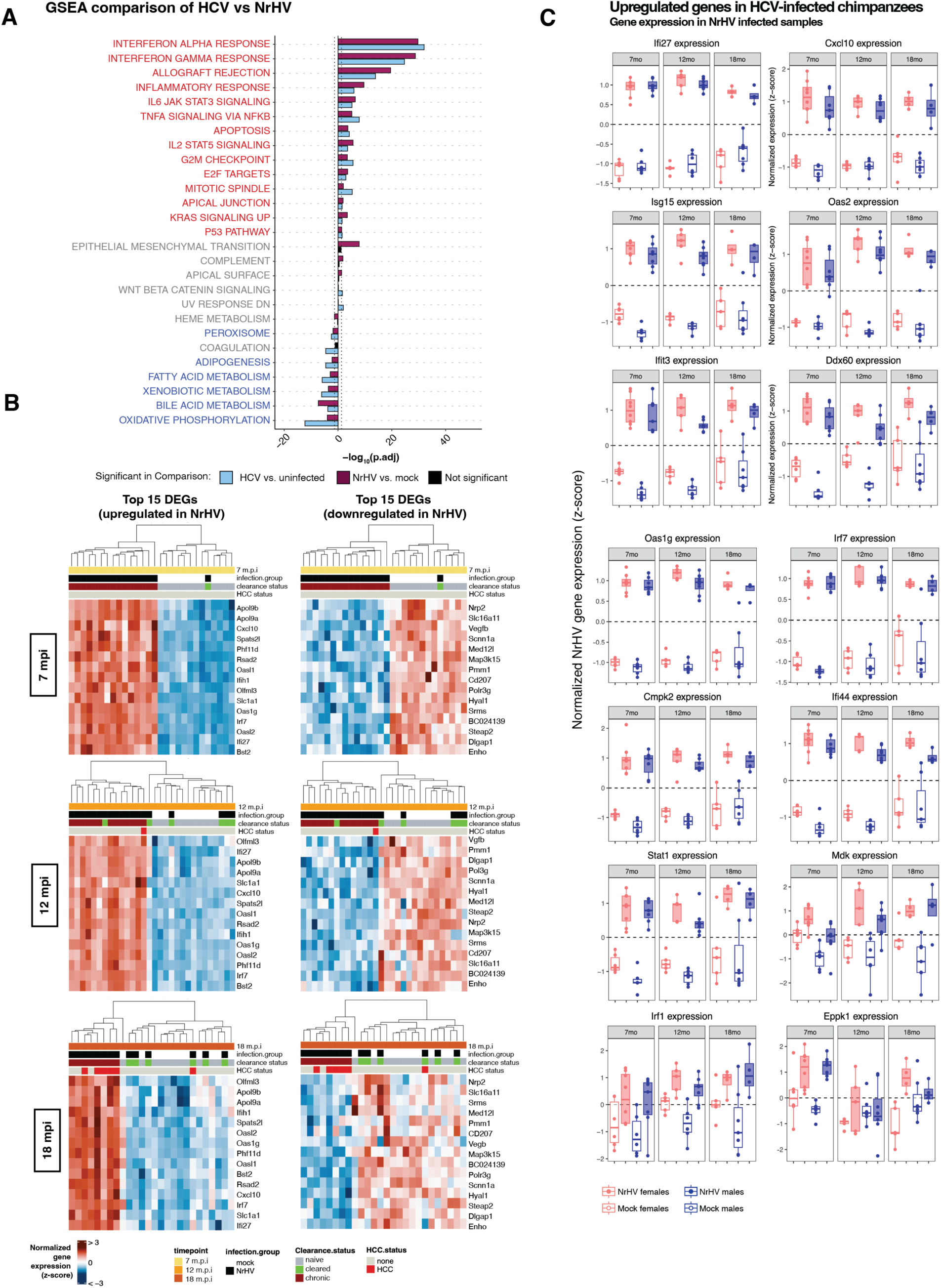
Differentially expressed genes (RNA-Seq) in NrHV vs mock-infected mice. (**A**) Gene Set Enrichment Analysis (GSEA) for HCV and NrHV (18 m.p.i.) showing HALLMARK gene sets enriched for each hepacivirus infection vs. respective controls. **(B)** Top 15 differentially expressed genes (up, left panel), ranked by adjusted p-value and filtered by log2 fold change > 1 in chronic NrHV infection, include multiple ISGs. Top 15 differentially expressed genes (down, right panel), ranked by adjusted p-value and filtered by log2 fold change < −1, include lipid metabolism genes and other genes associated with normal liver function, suggesting a loss of normal liver function. Time point (months post infection), Infection group (mock vs NrHV), Clearance status (naïve, cleared at any time point, chronic), HCC status (indicates if the mouse developed tumors – analysis is restricted to non-tumor tissue). (**C**) Normalized gene expression (z-scaled) of the 14 most upregulated genes identified from liver tissue from chronic HCV infection in chimpanzees(*57*) is displayed in NrHV-infected mice and age-matched mock controls over time. Salmon (females), dark blue (males), filled box plots (infected), unfilled box plots (mock). All samples are from cohort 2.

**S7.**
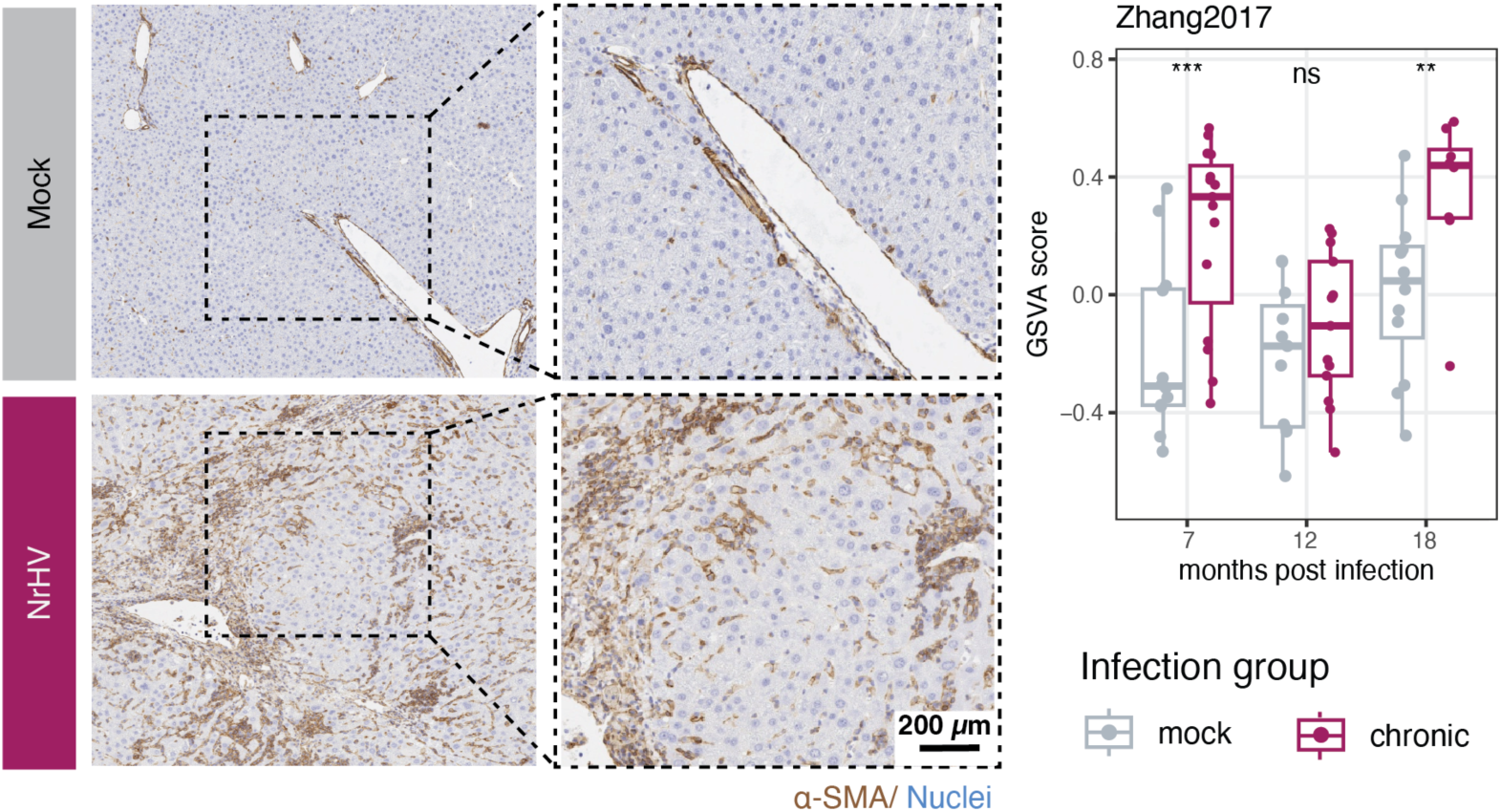
Hepatic Stellate Cell Signatures associated with cirrhosis and HCC. **(A)** Representative images of α-SMA in livers 18 m.p.i from mock- (gray) and NrHV- (maroon) infected livers. Enrichment of activated and quiescent hepatic stellate cell (HSC) gene set signature derived from Zhang et al., 2017(*27*) was analyzed in the context of mock or chronic NrHV infection. Each point represents one sample profiled by RNA-Seq, excluding tumor samples. A generalized linear model was used to calculate significance of enrichment across chronic and mock infection for each time point: adjusted P value < 0.001***; ns = not significant. All samples are from cohort 2.

**S8.**
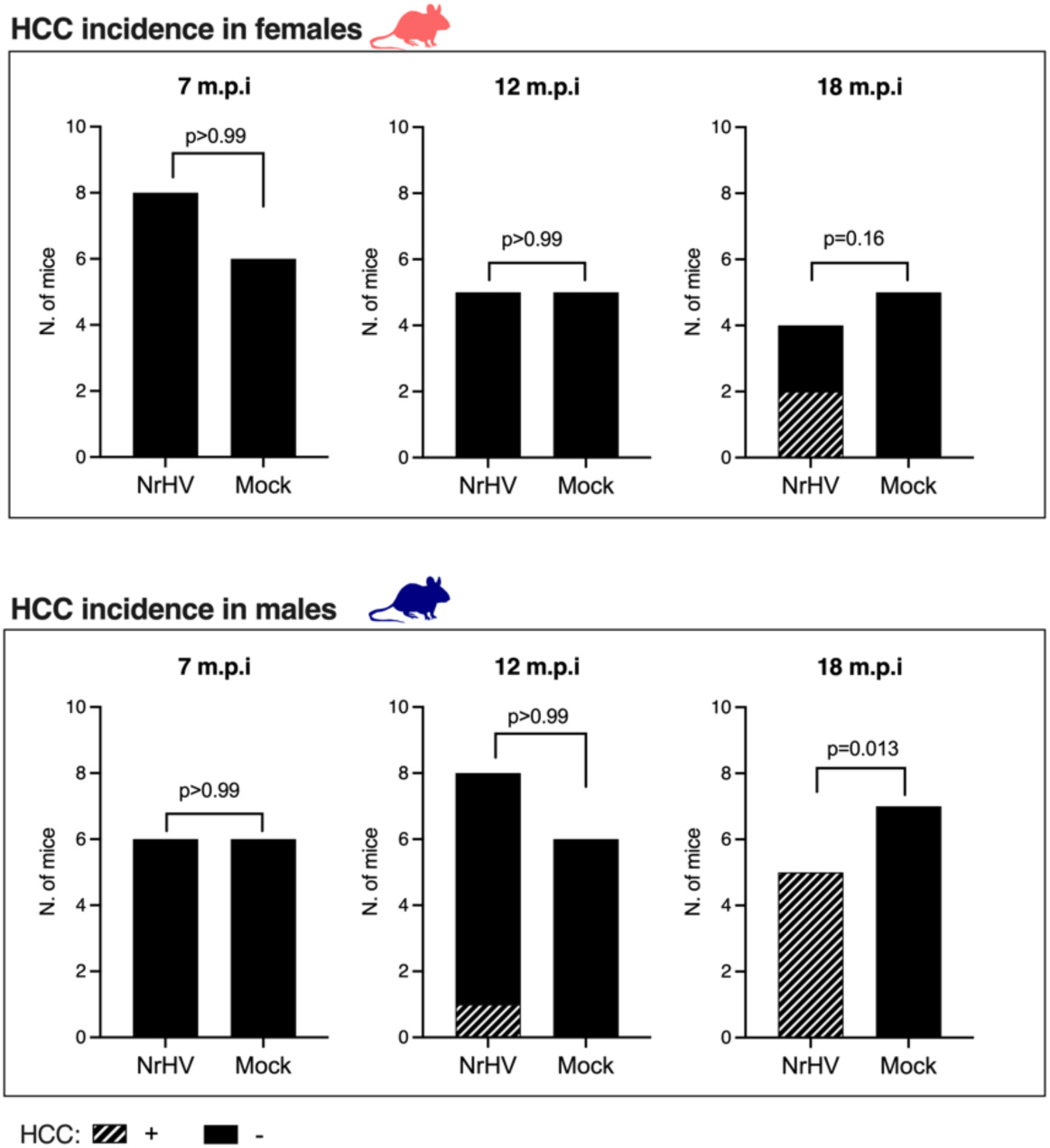
HCC frequency in NrHV-infected mice over time. Livers resected 7-, 12-, and 18-months post-infection were stained with H&E, blinded, and analyzed by two independent pathologists. Statistics: Fisher’s exact test (7 m.p.i: n=8 NrHV per sex, 6 mock per sex, ns: not significant/ 12 m.p.i: n=5 NrHV females, 6 NrHV males, 5 mock females, 6 mock males, ns: not significant/ 18 m.p.i: (n=4 NrHV females, 5 NrHV males, 5 mock females, 7 mock males). Mock controls did not present with tumors at any time point in cohort 2. All samples are from cohort 2 only.

**S9.**
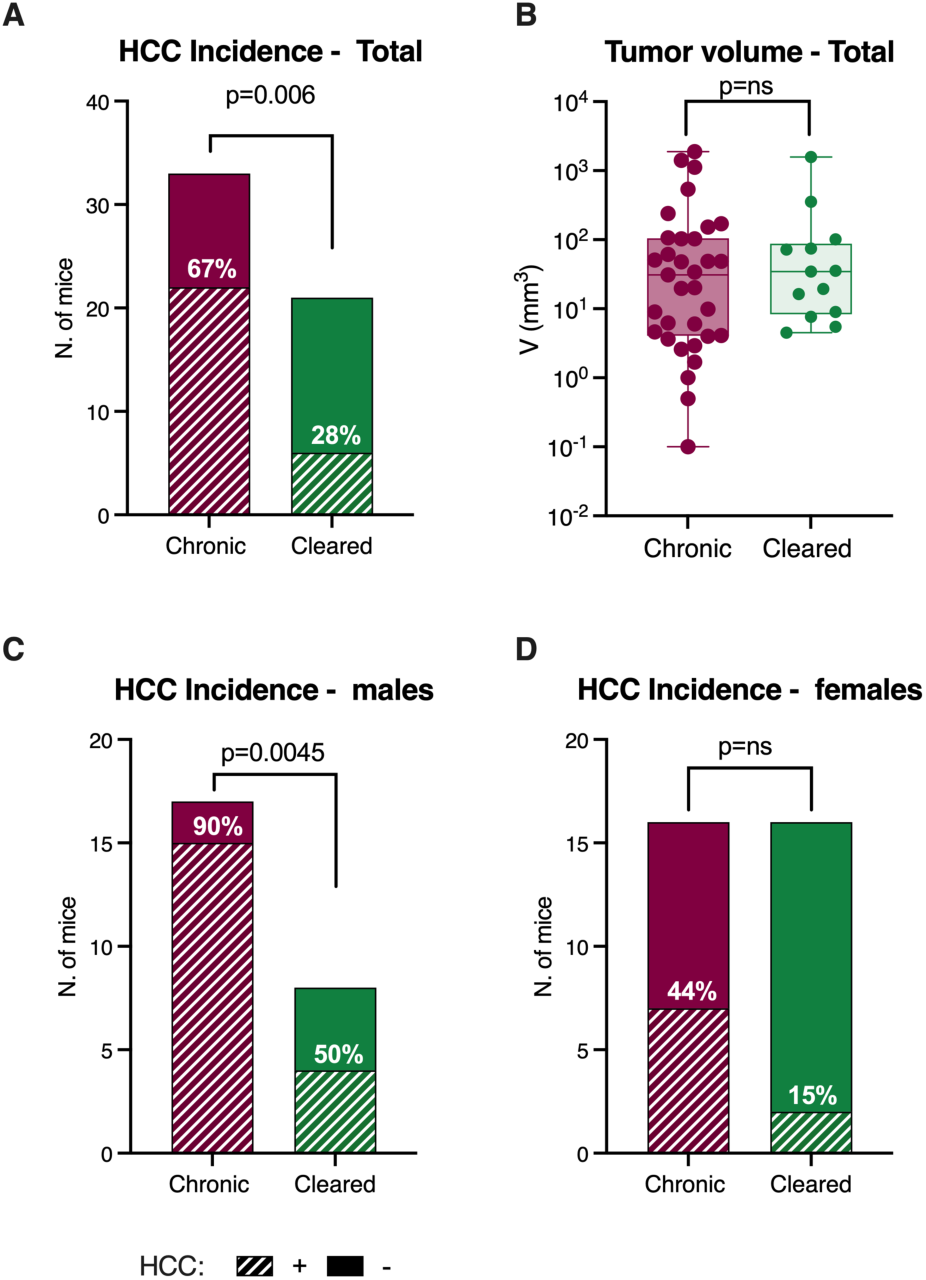
HCC development upon chronic vs resolved NrHV infection in mice. HCC incidence in NrHV-infected mice which cleared or not (chronic) the infection **(A)** Total HCC frequency in chronic vs cleared, **(B)** Volume of tumors in chronic vs cleared mice. Each dot represents an individual tumor, multiple tumors from a single mouse were included when present. **(C)** HCC incidence in males. **(D)** HCC incidence in females. Percentages on the bars indicate the frequency of mice with HCC (+HCC). Statistics: Fisher’s exact test (total: n=33 chronic, 21 cleared; males: n=17 chronic, 8 cleared; females: 16 chronic, 13 cleared). All spontaneously cleared mice alive at 18 m.p.i were included, irrespective of the time of clearance. These are combined data from two independent experiments collected 18 m.p.i (cohort 1 and 2).

**S10.**
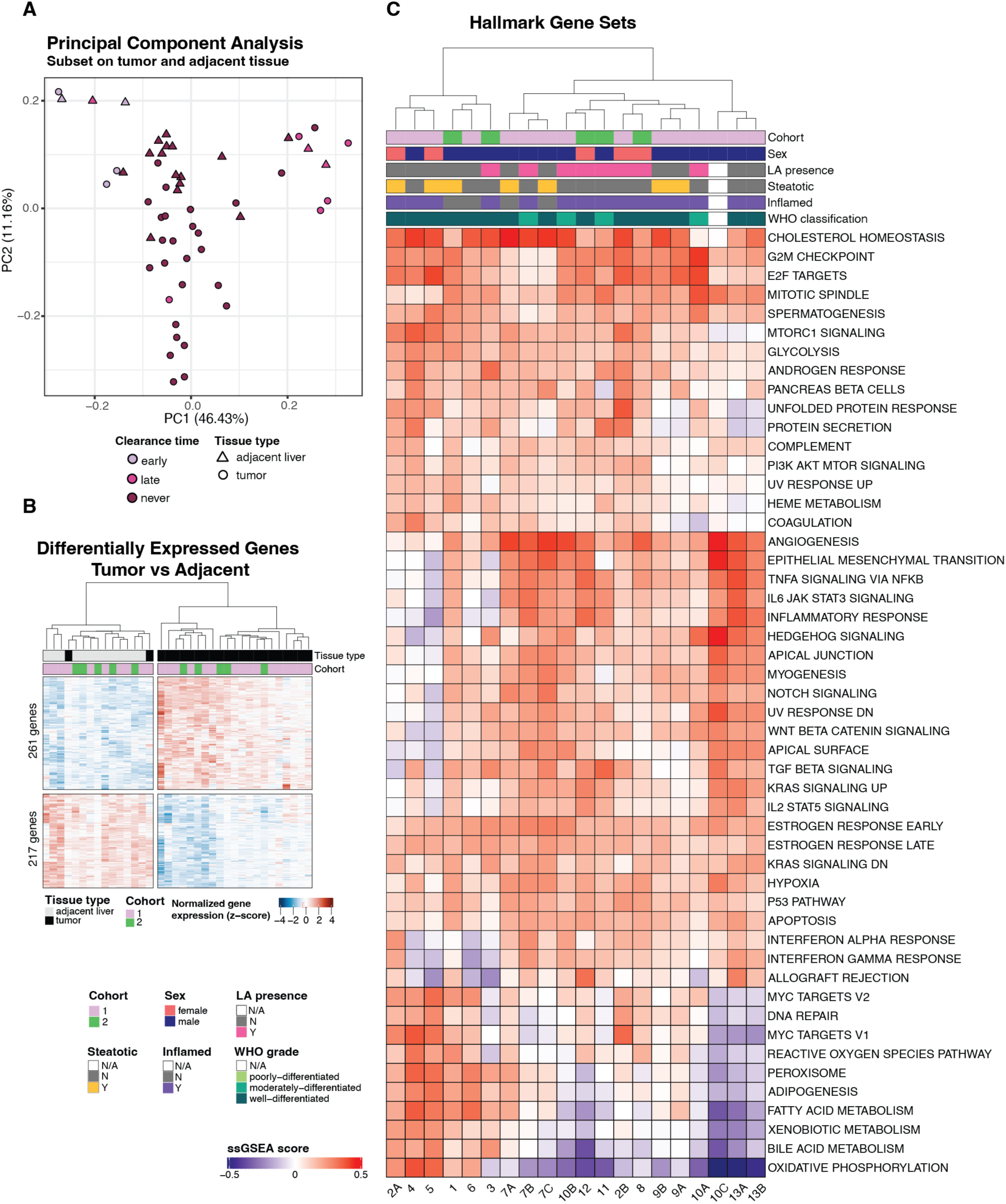
NrHV-elicited tumors are highly heterogeneous. **(A)** Principal component analysis of tumor and adjacent liver tissue from both NrHV infection cohorts. **(B)** Significantly highly and lowly expressed genes in tumor vs adjacent tissue across cohorts; significance: p.adj < 0.05 and absolute| log2FC | > 1. **(C)** Heatmap of all Hallmark gene sets showing ssGSEA scores for individual tumor and adjacent liver tissue pairs from RNA-Seq (n=20). All samples collected at 18 m.p.i (cohort 1 and 2, as labeled).

**S11.**
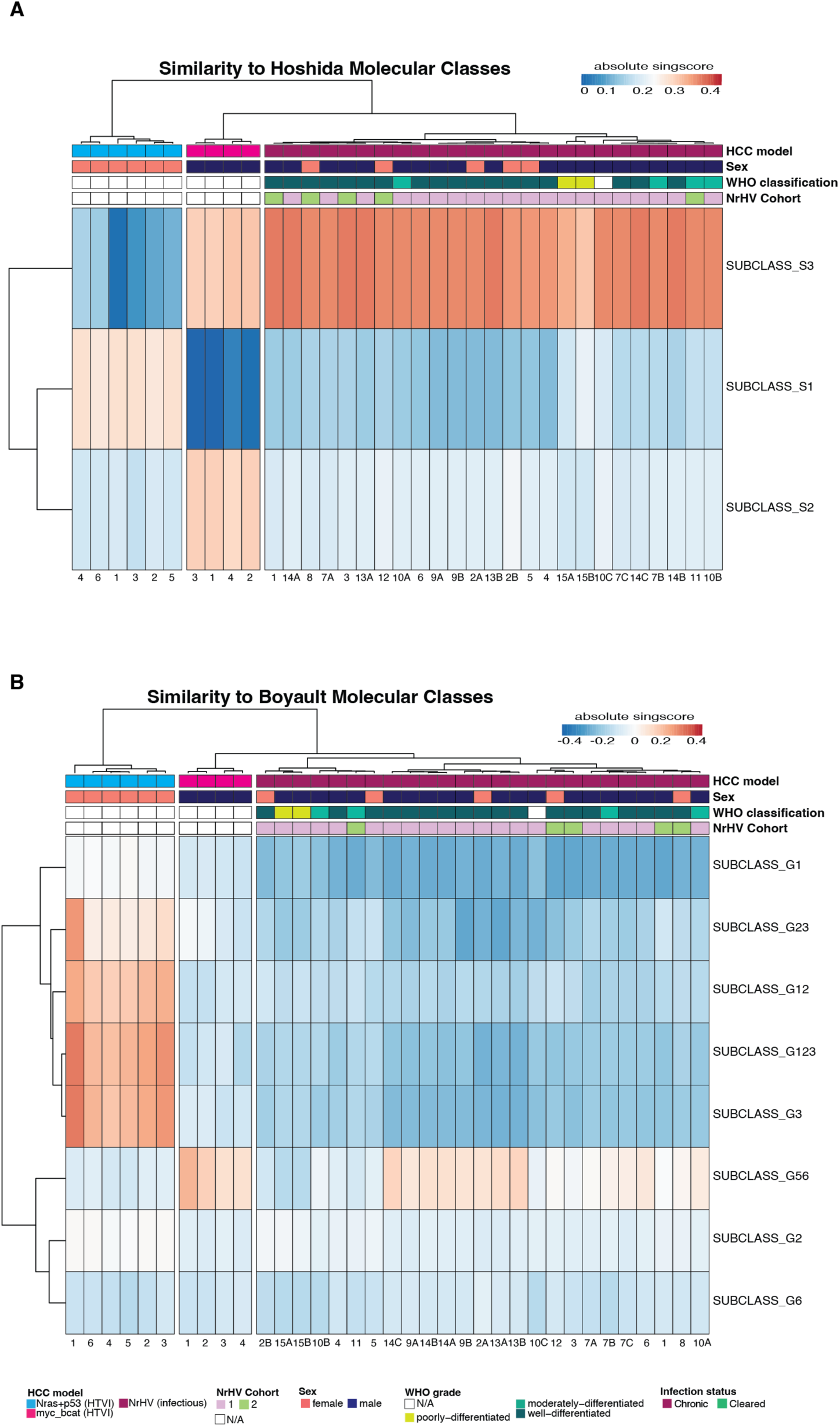
Similarity Scores (Singscore) of NrHV-elicited tumors compared to the Hoshida and Boyault human HCC molecular subclasses. **(A)** Hoshida. **(B)** Boyault. All tumors from NrHV-infected mice which had high quality RNA were included in this analysis irrespective of adjacent pairing (n=25 tumors, from 15 mice). HTVI samples from ≈-catenin(*37*) and p53(*36*) mutant tumor models were used to highlight differences among different tumor types p53 represents the “proliferation” and ≈-catenin represents the “non-proliferation” cholestatic classes of human HCC respectively.

**S12.**
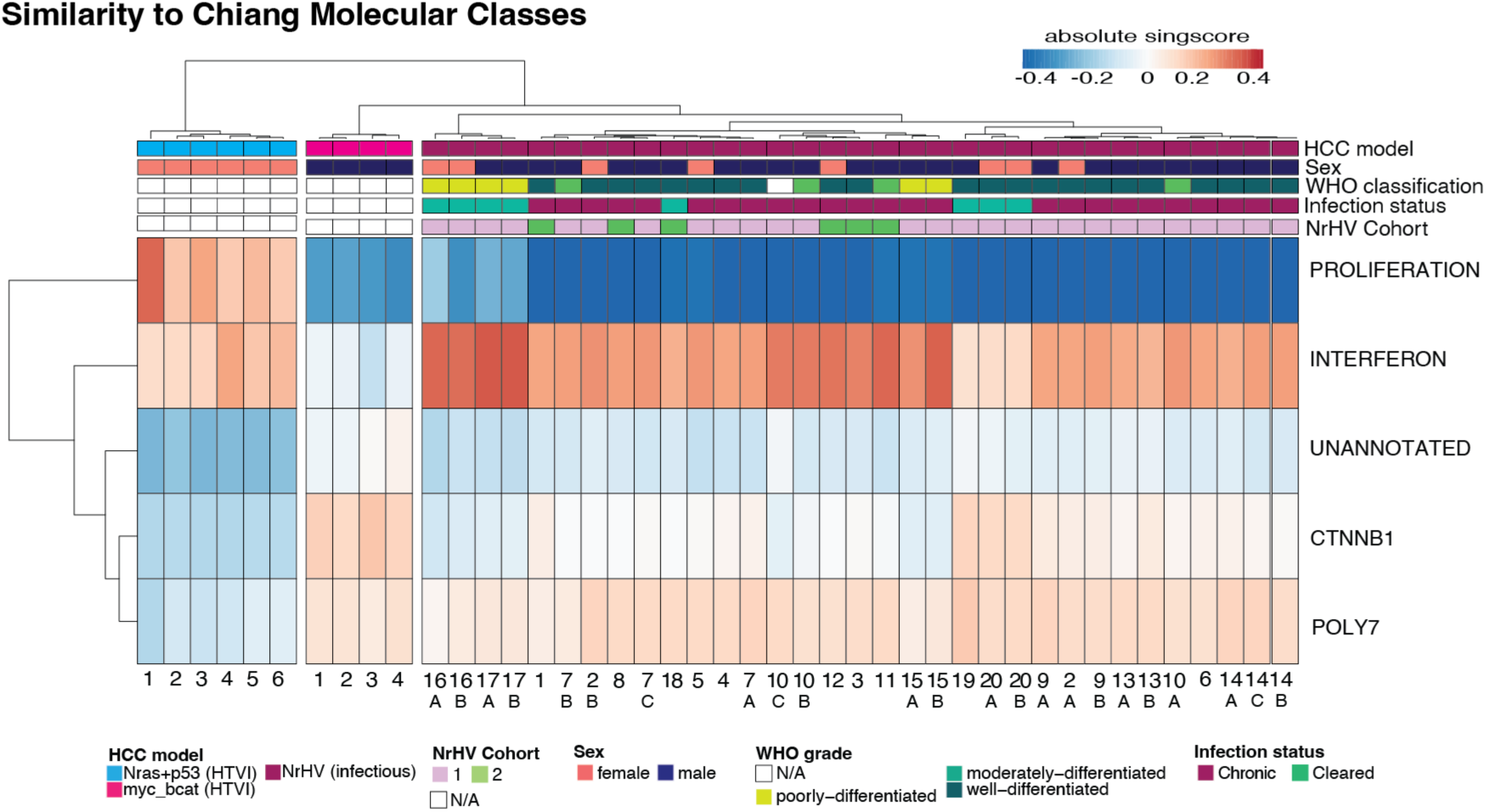
Tumors in mice that spontaneously cleared virus are transcriptionally distinct from tumors from chronically infected mice. Singscore comparing similarity between cleared and chronic tumors and the Chiang HCC molecular classification of human HCV-elicited tumors. Total number of cleared mice: 22; cleared mice with tumors: 6; total number of tumors in the cleared group: 12. RNA-sequenced tumors: cleared (n=8) vs. chronic (n=26). Tumors occurring in the same mouse are represented by letters. Data from samples collected at 18 m.p.i (cohort 1 and 2, as labeled). HTVI samples from ≈-catenin(*37*) and p53(*36*) mutant tumor models were used to highlight differences among different tumor types p53 represents the “proliferation” and ≈-catenin represents the “non-proliferation” cholestatic classes of human HCC respectively.

**Table S1.** Wild-type C57BL/6J mouse cohorts summary.

| **Cohort 1** |  | **Harvest 18 m.p.i** |  |
| --- | --- | --- | --- |
|  |  | *Liver* | *HCC* |
| **NrHV females** |  |  |  |
| Total Launched | 30 |  |  |
| Chronic infection | 29 | 12 | 5 |
| Spontaneous clearance | 9 | 9 | 2 |
| Mortality | 8 |  |  |
| **Total harvested females** | **21** | 21 | 7* |
| **NrHV males** |  |  |  |
| Total Launched | 30 |  |  |
| Chronic infection | 29 | 12 | 10 |
| Spontaneous clearance | 7 | 7 | 3 |
| Mortality | 10 |  |  |
| **Total harvested males** | **19** | **19** | 13* |
| **Mock-infected females** |  |  |  |
| Launched | 10 | 3 | 0 |
| Mortality | 2 |  |  |
| **Total harvested mock female** | **3\*\*** |  |  |
| **Mock-infected males** |  |  |  |
| Launched | 10 | 10 | 1 |
| Mortality | 0 |  |  |
| **Total harvested mock male** | **10** |  |  |
| **Total launched** | **80** |  |  |

*Note: \*mice with multiple tumors were counted only once; \*\* one cage of mock females (n=5) was found flooded and these mice were excluded from all analysis. Chronic refers to infection detected $\geq 1$ month post-infection. Mice with no infection detected within the first month were excluded (failed infection) and were not accounted later in the cleared group.*

|  |  | Harvest 7 m.p.i |  | Harvest 12 m.p.i |  | Harvest 18 m.p.i |  |
| --- | --- | --- | --- | --- | --- | --- | --- |
| Cohort 2 |  | Liver | HCC | Liver | HCC | Liver | HCC |
| NrHV females |  |  |  |  |  |  |  |
| Total Launched | 30 |  |  |  |  |  |  |
| Chronic infection | 29 | 7 | 0 | 5 | 0 | 4 | 2 |
| Spontaneous clearance | - | 0 | 0 | 5 | 0 | 4 | 0 |
| Mortality | 3 |  |  |  |  |  |  |
| **Total harvested** | **26** | **7** | **0** | **10** | **0** | **8** | **2** |
| NrHV males |  |  |  |  |  |  |  |
| Total Launched | 30 |  |  |  |  |  |  |
| Chronic infection | 29 | 7 | 0 | 8 | 1 | 5 | 5 |
| Spontaneous clearance | - | 0 | 0 | 1 | 0 | 1 | 1 |
| Mortality | 7 |  |  |  |  |  |  |
| **Total harvested** | **22** | **7** | **0** | **9** | **1** | **6** | **6** |
| Mock-infected females |  |  |  |  |  |  |  |
| Launched | 16 | 6 | 0 | 5 | 0 | 5 | 0 |
| Mortality | 0 |  |  |  |  |  |  |
| **Total harvested mock female** | **16** |  |  |  |  |  |  |
| Mock-infected males |  |  |  |  |  |  |  |
| Launched | 20 | 6 | 0 | 6 | 0 | 6 | 0 |
| Mortality | 2 |  |  |  |  |  |  |
| **Total harvested mock male** | **18** |  |  |  |  |  |  |
| **Total launched** | **96** |  |  |  |  |  |  |

*Note: one uninfected male developed a spleen tumor noted at the termination of the experiment and was excluded from all analyses. Chronic refers to infection detected $\geq 1$ month post-infection. Mice with no infection detected within the first month were excluded (failed infection) and were not accounted later in the cleared group.*

**Table S2.**
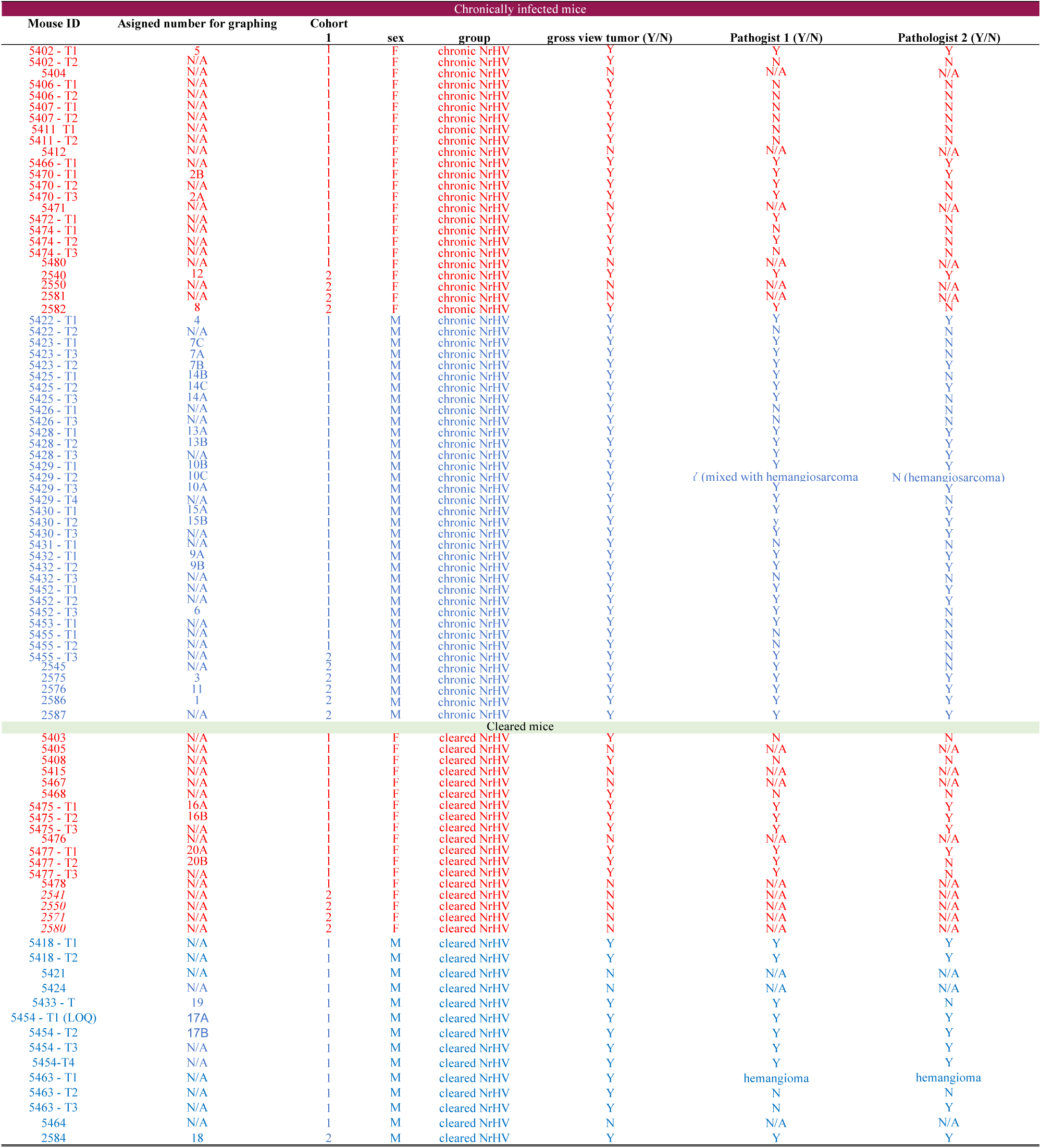
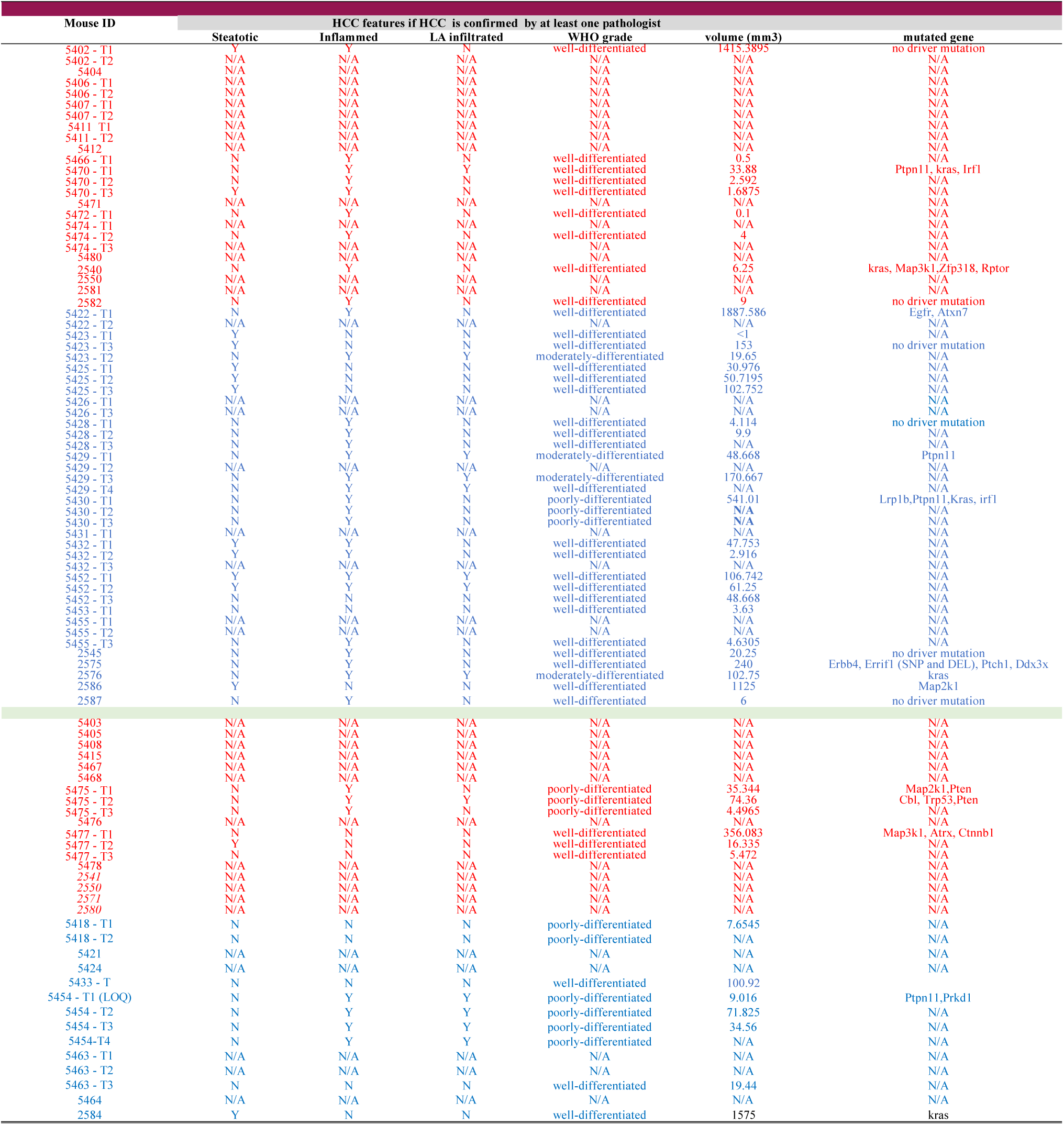
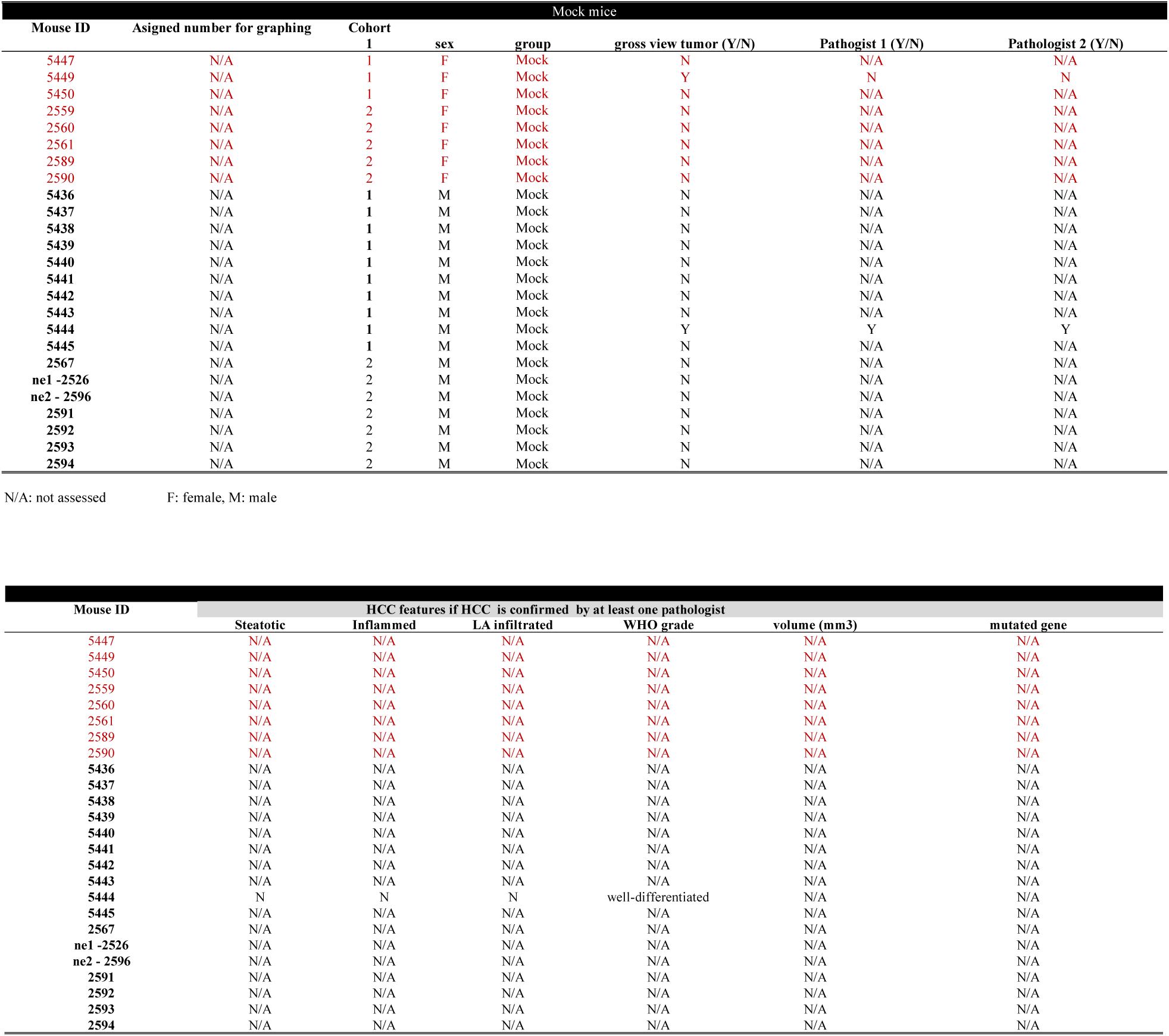
Tumor general features.

**Table S3.** Fibrosis and Inflammation Score Criteria.

| **Mouse Fibrosis Score** | **Description** |  |
| --- | --- | --- |
| **0** | No fibrosis |  |
| **1** | Fibrous expansion of some portal areas |  |
| **2** | Fibrous expansion of most portal areas |  |
| **3** | Fibrous expansion of most portal areas with occasional portal to portal (P-P) bridging |  |
| **4** | Fibrous expansion of portal areas with marked bridging (P-P) and portal-central (P-C) |  |
| **5** | Marked bridging (P-P and/ or P-C) with occasional nodules – transition to cirrhosis |  |
| **6** | Established cirrhosis |  |
| **Mouse Inflammation score (Activity)** | **Interface Hepatitis** | **Parenchymal Injury** |
| A0 | None | Less than one focus per lobule |
| A1 | None | At least one per lobule |
|  | Focal, some portal areas | Less than several per lobule |
| A2 | None | Several per lobule |
|  | Focal, some portal areas | Several per lobule |
|  | Focal, all, or diffuse, some portal areas | Less than several per lobule |
| A3 | Focal, all, or diffuse, some portal areas | Several per lobule |
|  | Diffuse, all portal areas | Any amount |

*Note: criteria for Activity are based on acidophil bodies and immune infiltration as described in Goodman, Journal of Hepatology, 2007(22)*

**Table S4.**
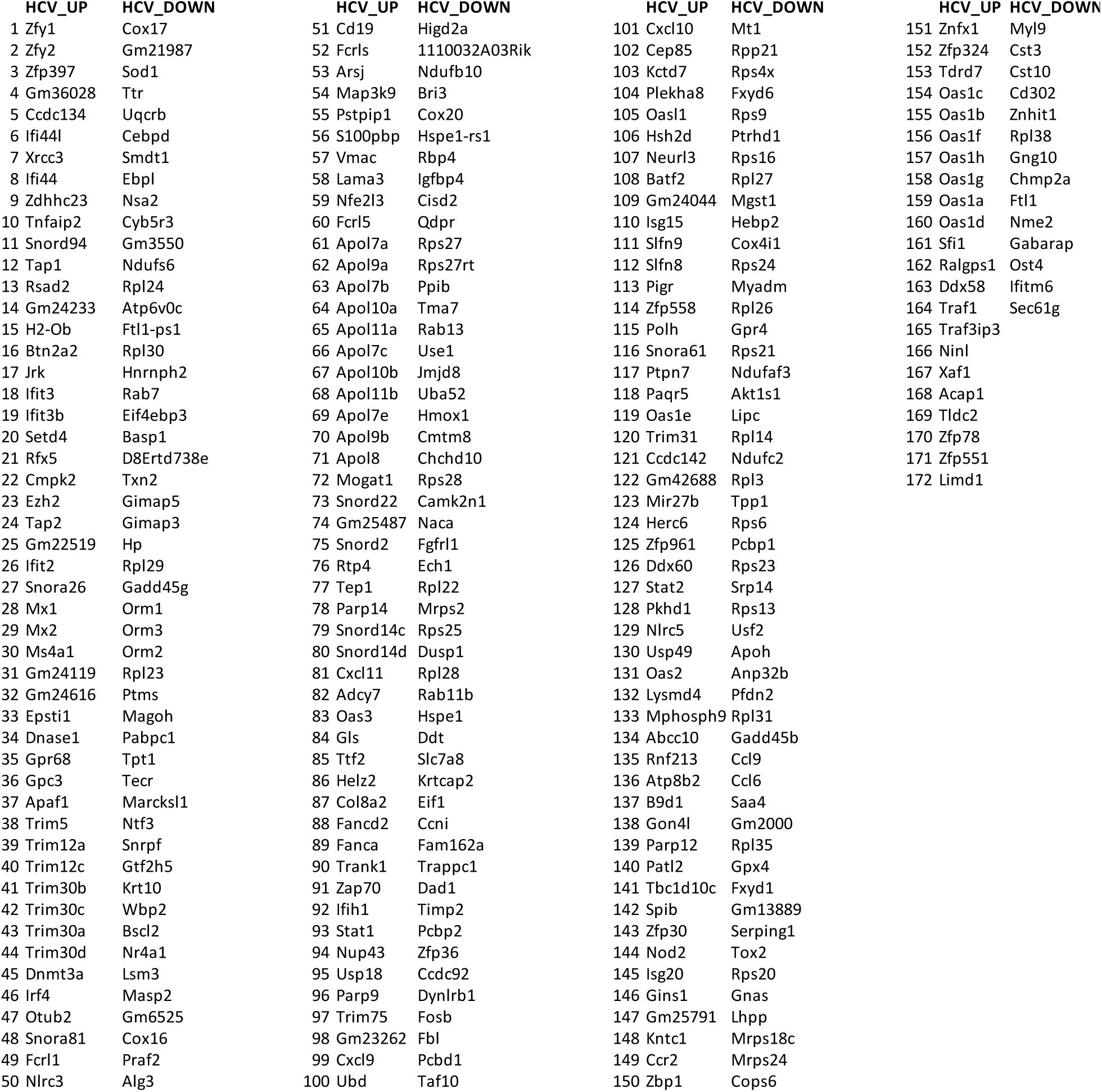
HCV Up and Down-regulated gene sets.

**Table S5.**
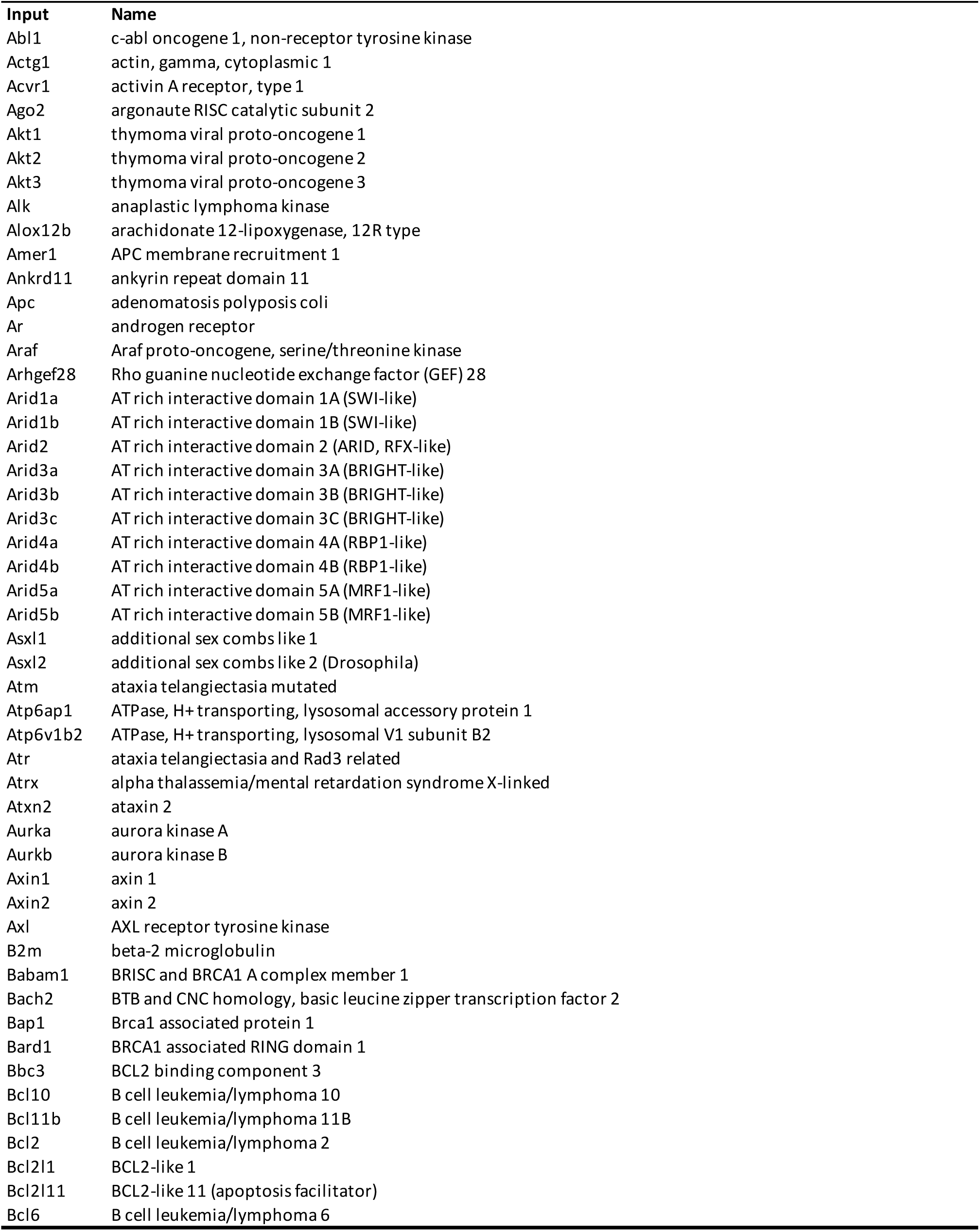
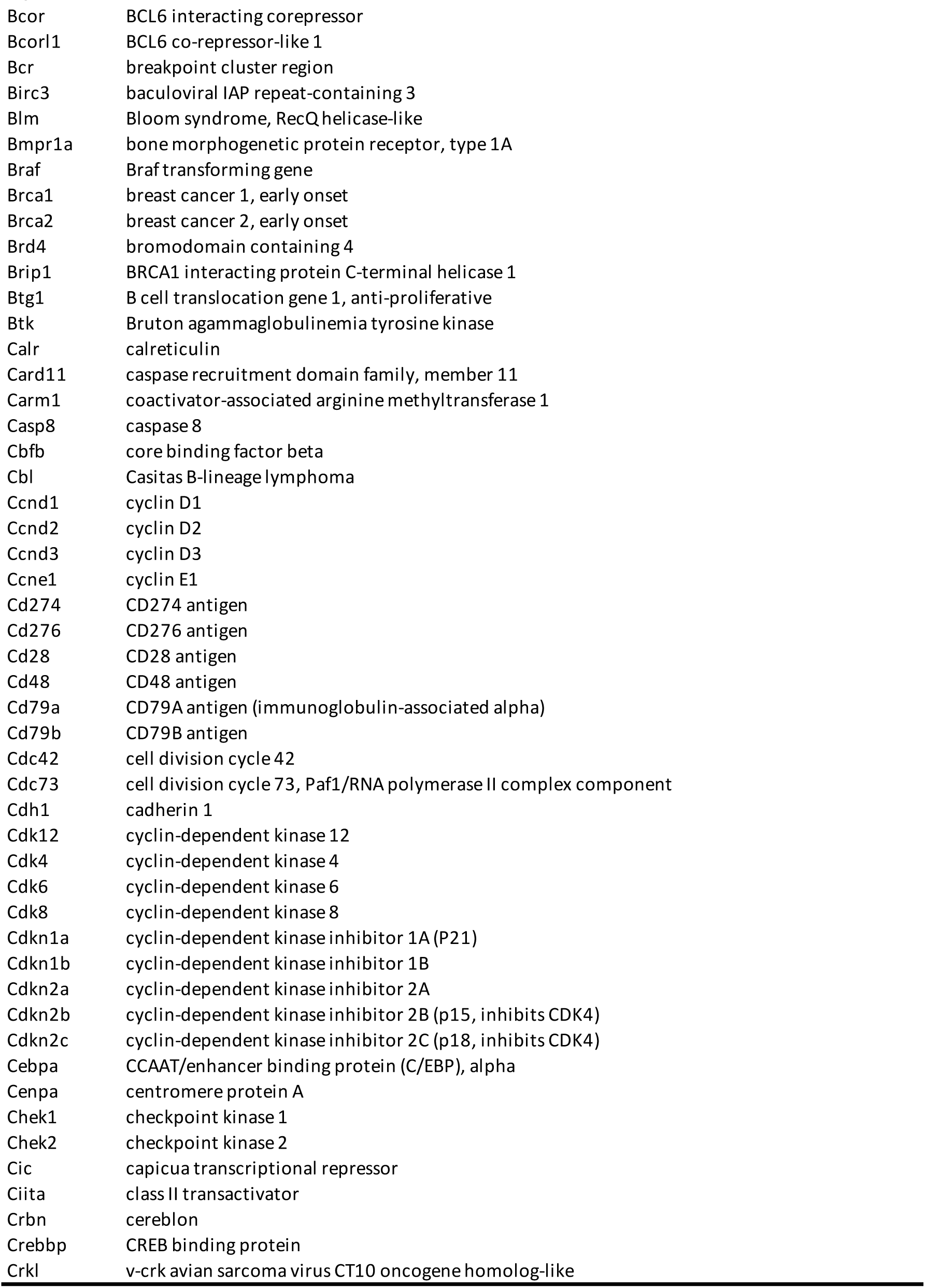
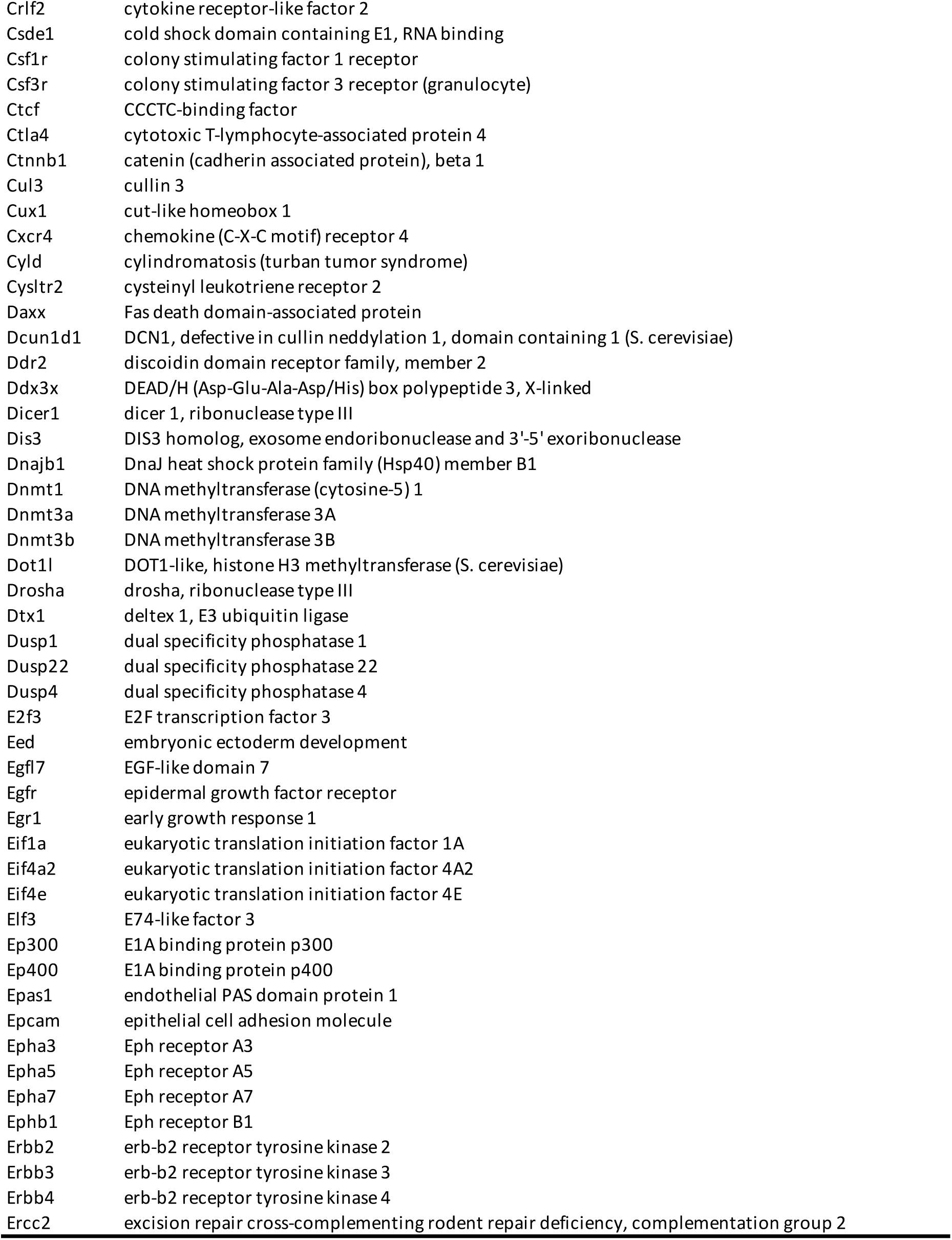
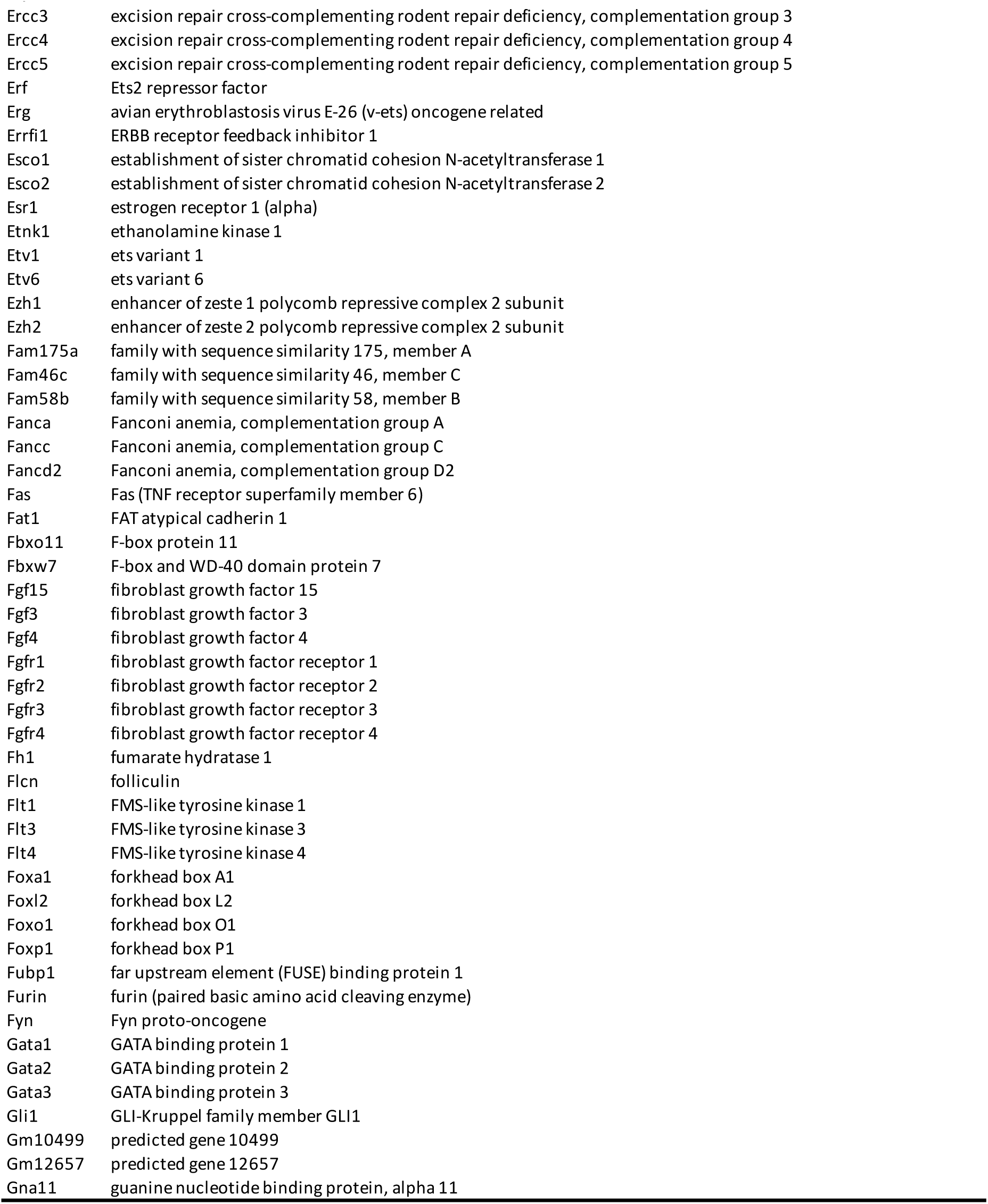
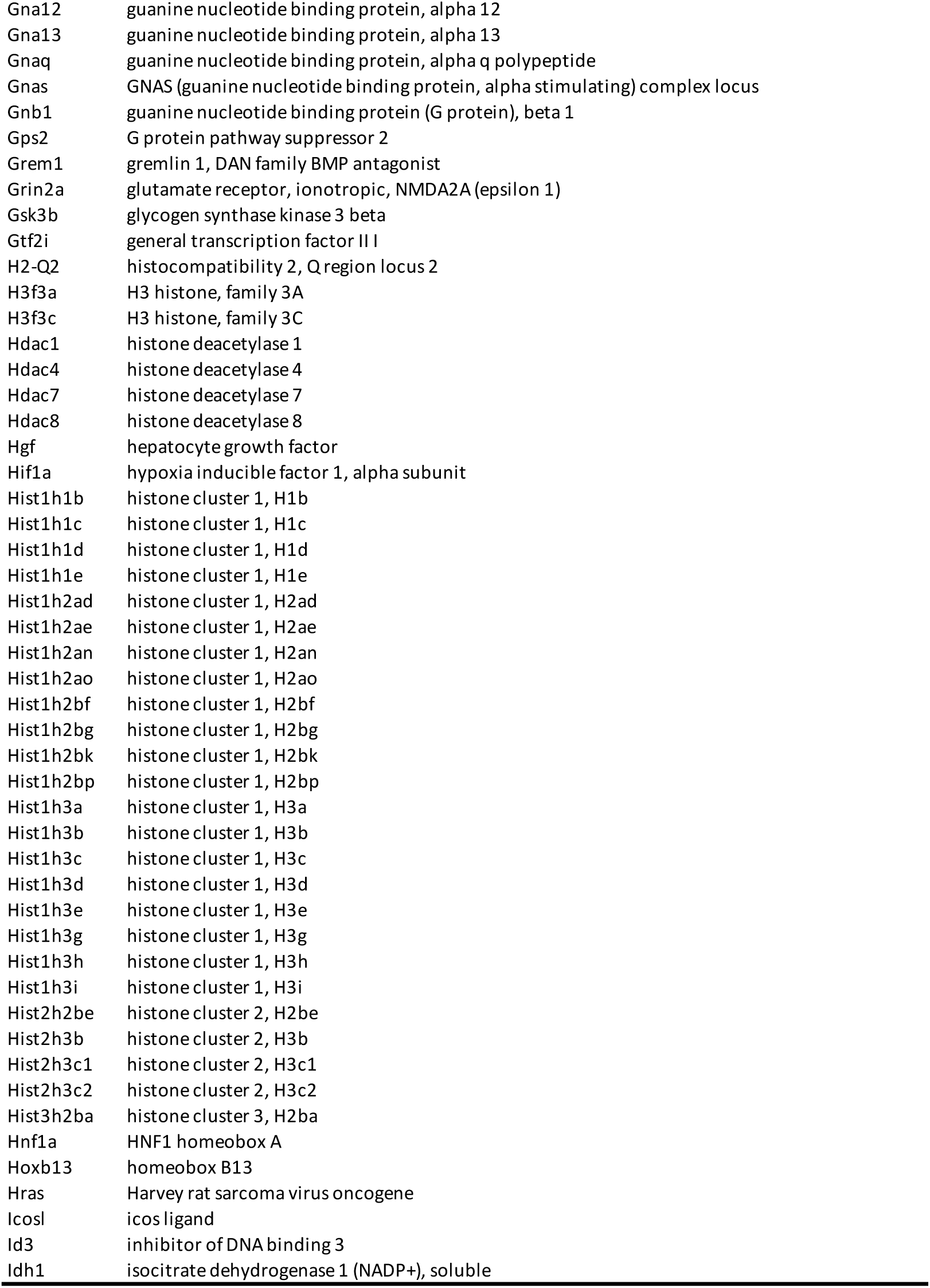
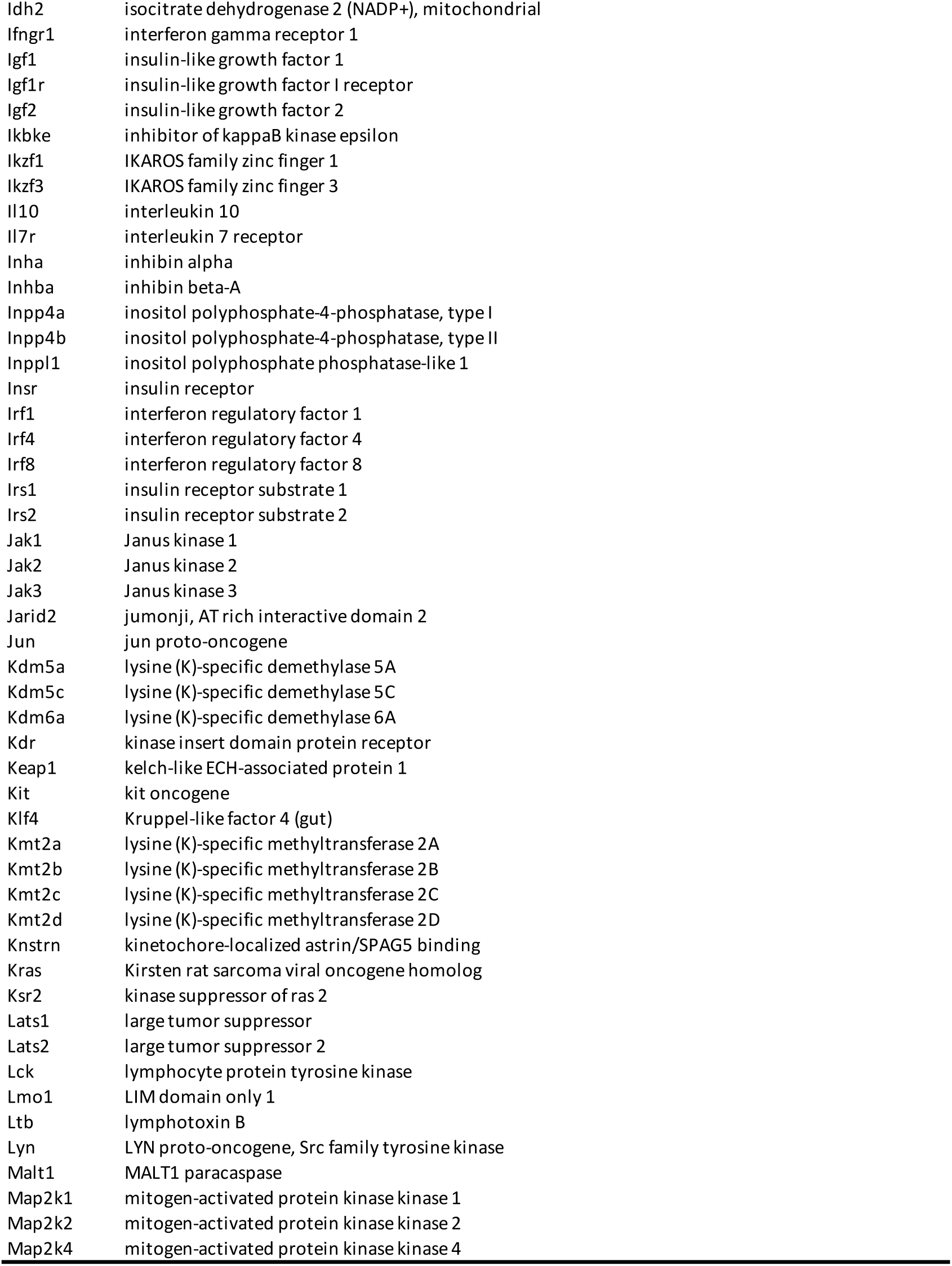
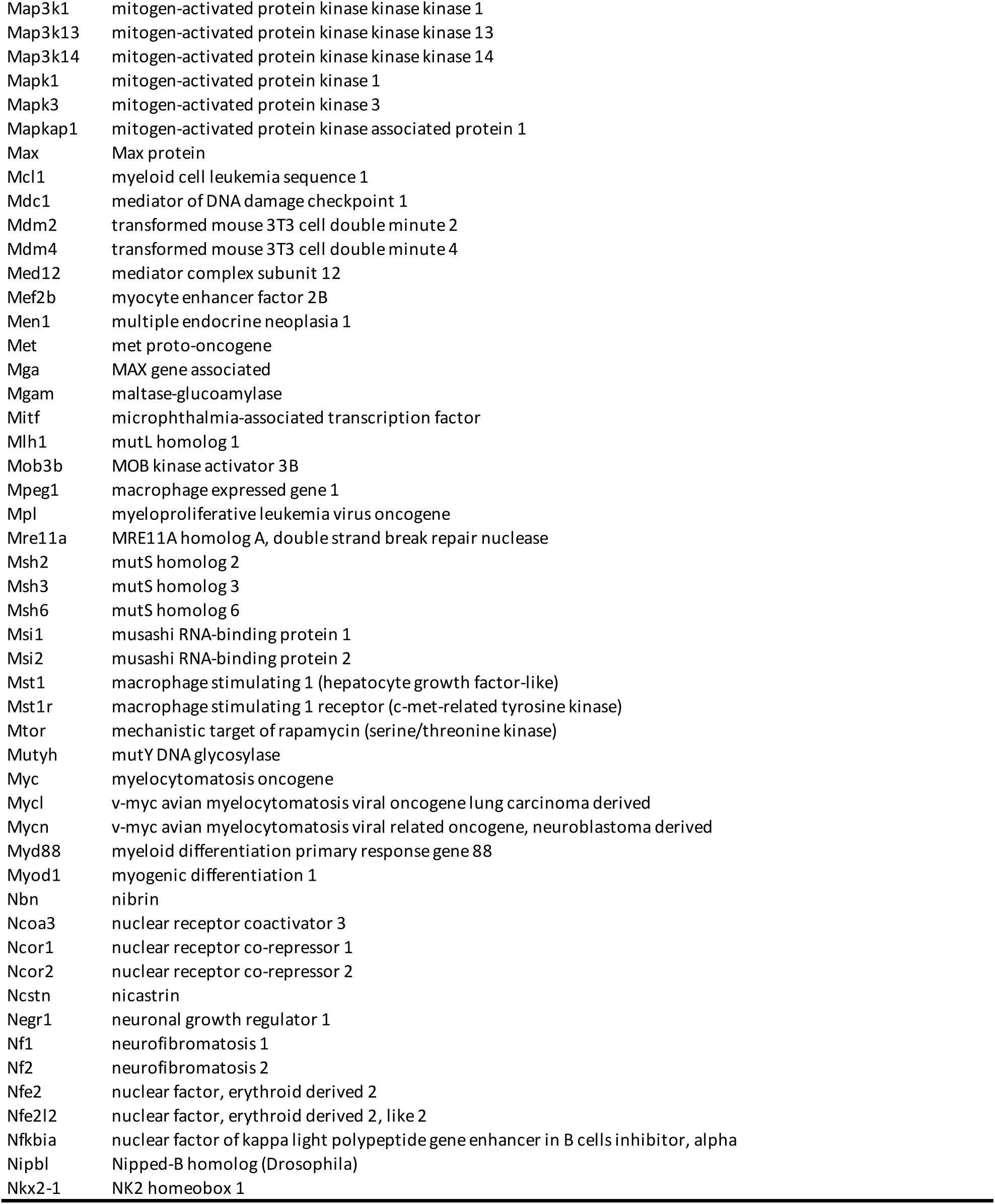
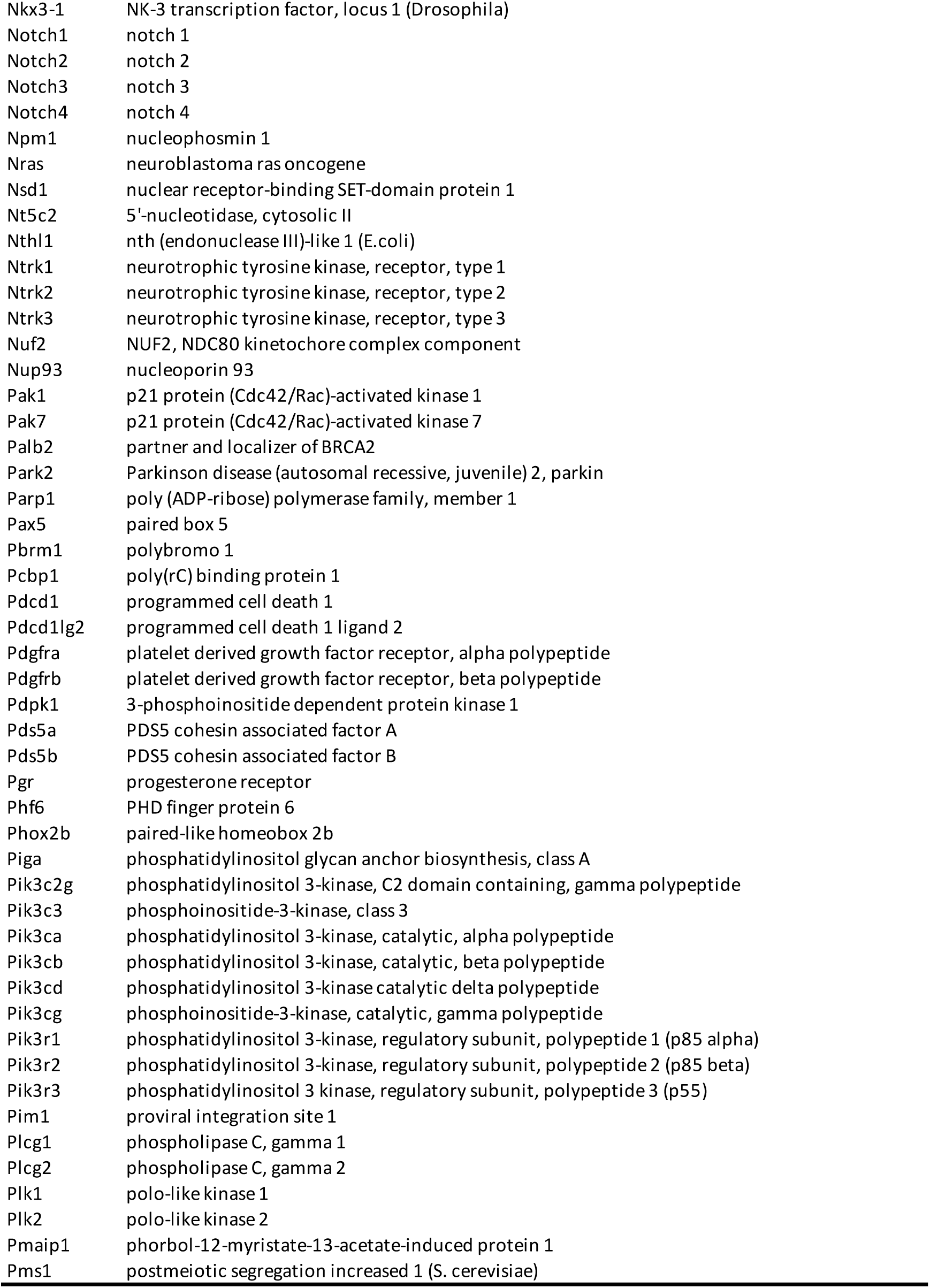
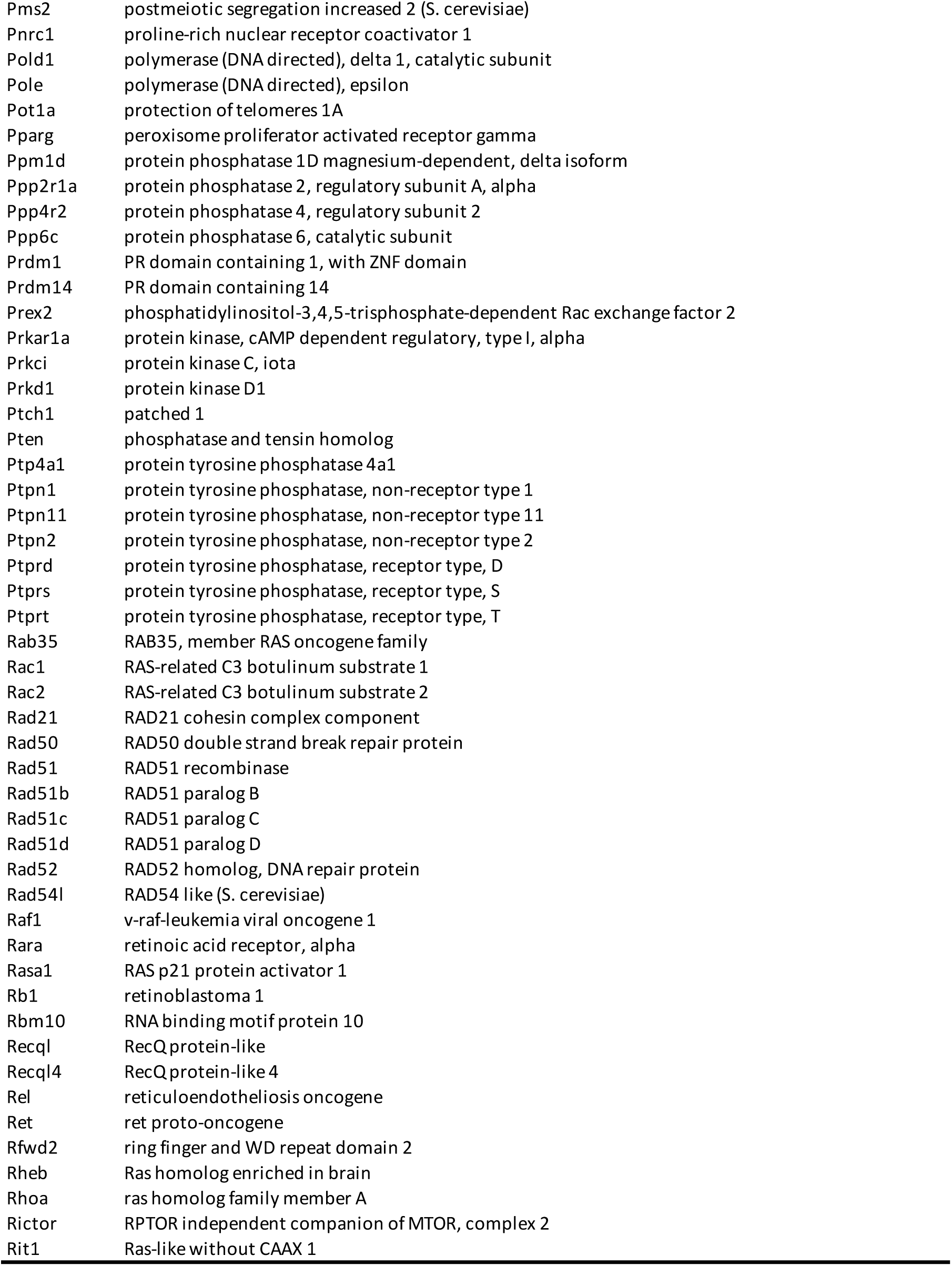
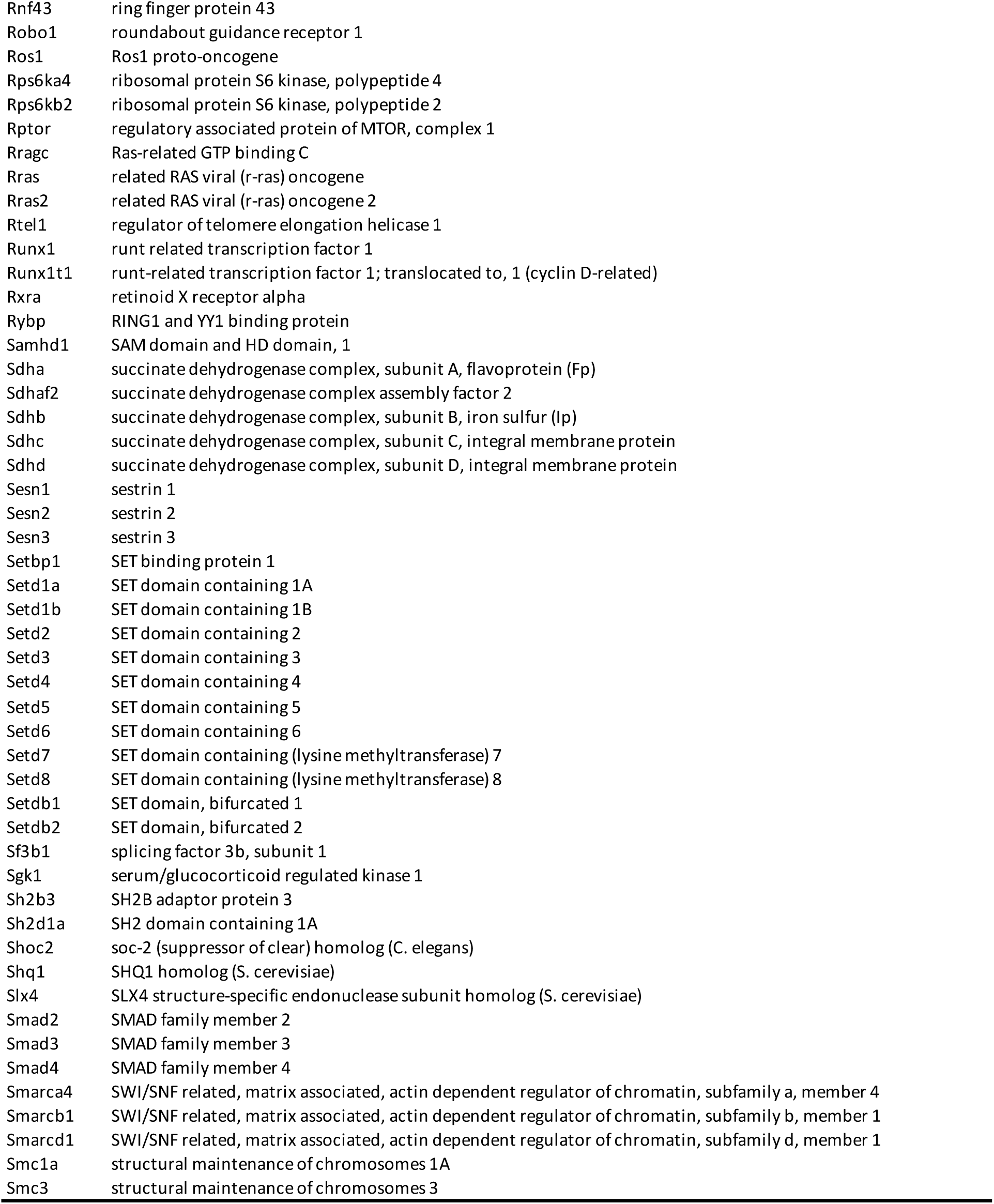
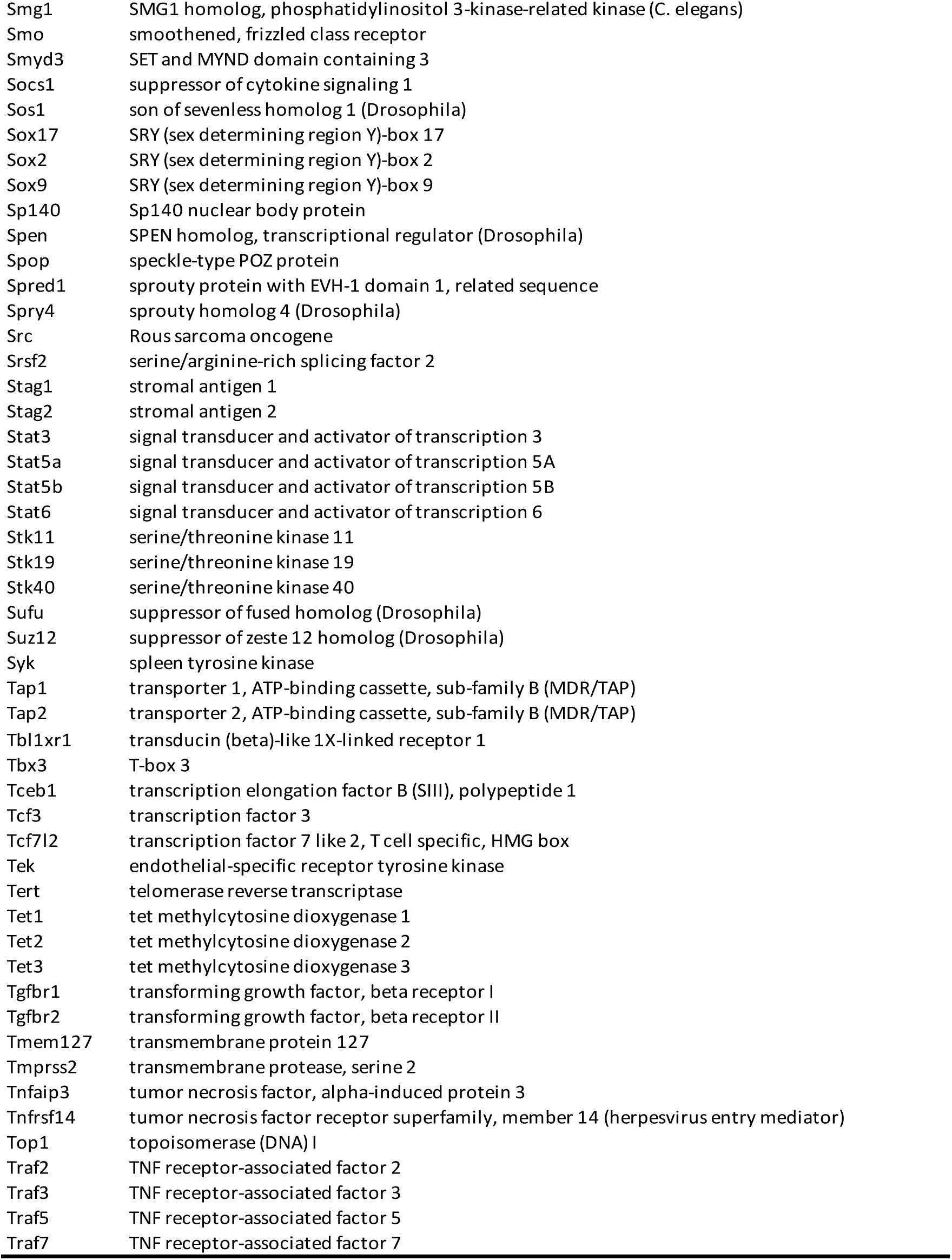
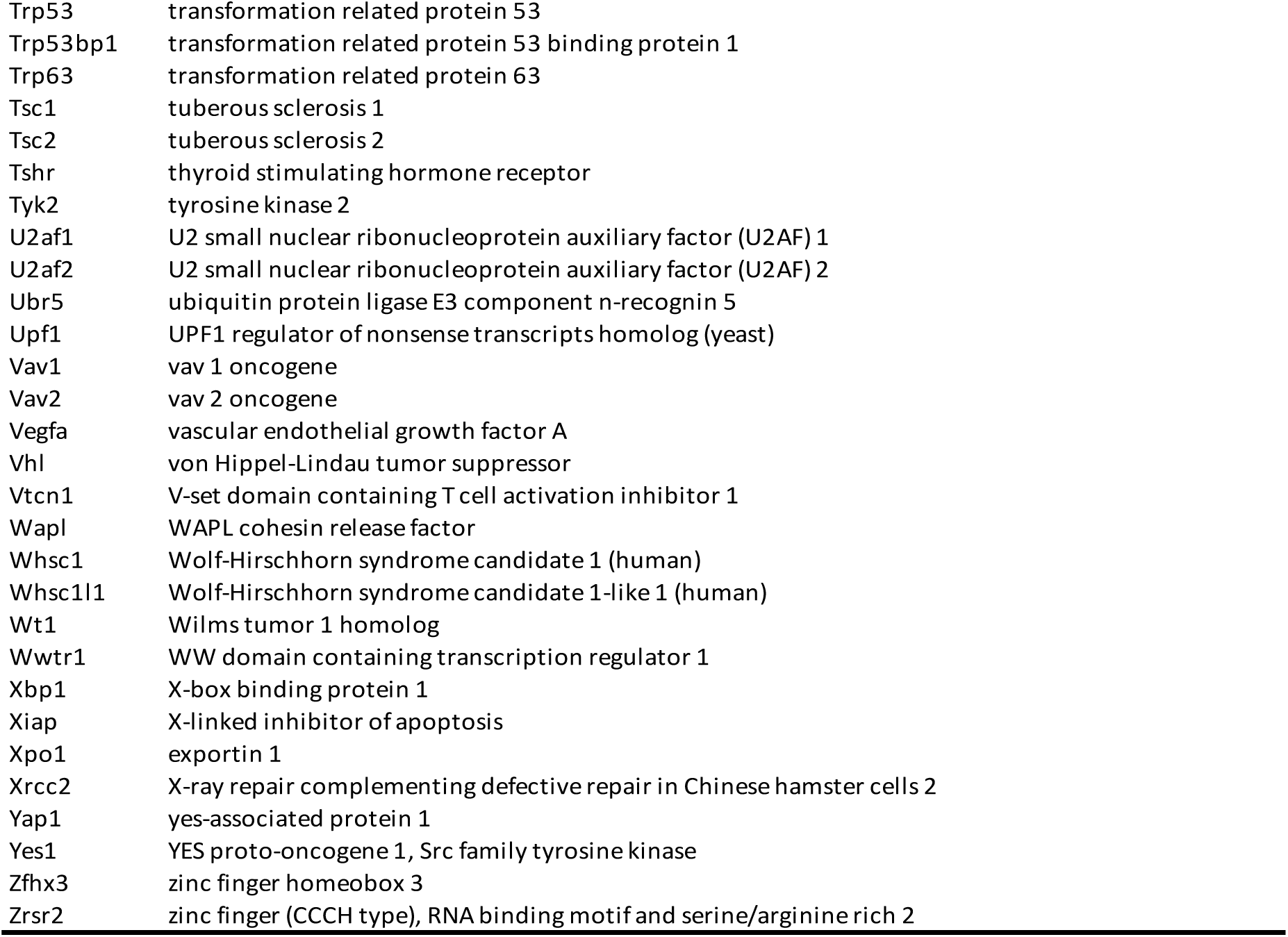
Genes/genic regions targeted in Mouse-IMPACT.

## Acknowledgments

We thank Dr. Scott Friedman, Dr. Andrea Schietinger and Dr. Utpal B. Pajvani for insightful discussions and for technical assistance in experimental design. We thank Dr. William Schneider, Dr. Lauren Aguado and Georgia McClain for their valuable feedback and editing. We acknowledge the Flow Cytometry Facility at the Rockefeller University for providing technical advice and the Genomics Core at Memorial Sloan-Kettering Cancer Center for DNA sequencing and data analysis.

## Funding

Stavros Niarchos Foundation (SNF) as part of its grant to the SNF Institute for Global Infectious Disease Research at The Rockefeller University (MNB, HAC, CMR)

This research was partially supported by the Marlene Hess Center for Research on Women’s Health and Biomedicine as part of its grant program at The Rockefeller University (MNB)

National Institutes of Health, grant R01 AI131688-05 (MNB, EB, BRR, CMR)

National Center for Advancing Translational Sciences (NCATS), National Institutes of Health (NIH) Clinical and Translational Science Award (CTSA) program, grant #UL1 TR001866 (CJ)

The European Research Council (ERC), Starting Grant 802899 (TKHS)

The Danish Cancer Society, grant R374-A22411 (TKHS).

FIOCRUZ and CNPq 312671-2022-2, 205096-2018-2 (JB)

FIOCRUZ and CAPES 88881.337176/2019-01 (ALPM)

## Author contributions

Conceptualization: MNB, CMR. Methodology: MNB, CMR, BRR. Investigation: MNB, JB, ALPM, TB, HAC, GTX, NP, ABV, TL, KHD, BZ, CQ, RW, CJ, BC, LC, NT. Visualization: MNB, TB, HAC, YJH, BRR. Funding acquisition: CMR, BRR, EB, TKHS, TPS, MNB. Writing – original draft: MNB, TB. Writing – review & editing: MRM, AK, TKHS, TPS, EB, SL, BRR, CMR

## Diversity, equity, ethics, and inclusion [optional]

The authors are committed to fostering diversity, equity, and inclusion in scientific research. This study was conducted in an environment that values equal opportunity and ethical research practices. We recognize the contributions of researchers from diverse backgrounds and strive to create an inclusive and supportive scientific community. Our work adheres to the highest ethical standards, including responsible data collection, analysis, and reporting. Additionally, we acknowledge the importance of equitable access to scientific knowledge and welcome collaboration across disciplines and institutions to advance inclusive research practices.

## Data and materials availability

All data supporting the findings of this study are available in the main text, supplementary materials or for download at the NCBI GEO repository under accession GSE265927.

## Supplementary Materials

S2, S4 and S5 provided as excel files

